# Multimodal sensory overload in dopamine-deficient larval zebrafish leads to paradoxical kinesia

**DOI:** 10.1101/2025.06.30.662454

**Authors:** Krishnashish Bose, Karnika Bhardwaj, Su Guo

## Abstract

Paradoxical kinesia—the temporary alleviation of motor deficits by powerful, urgent stimuli in Parkinson’s disease (PD)—remains poorly understood at the neural circuit level. Through chemo-genetic ablation of tyrosine hydroxylase-expressing neurons in larval zebrafish and brain-wide calcium imaging under head-fixed, tail-free conditions, we uncovered a neural mechanism underlying this phenomenon. While catecholamine (CA)-deficient larvae exhibited severe locomotor deficits during free swimming, they showed paradoxical recovery of tail movements during whole-brain neural activity imaging. This locomotor recovery was accompanied by a significantly increased number of active neurons in the midbrain and hindbrain, but with reduced firing rates. Further analyses across 2158 anatomically defined regions allowed us to uncover a subset of regions, genes, and neurotransmitter types. GABAergic neurons were found to primarily account for the hyperactivity in the hindbrain, while glutamatergic neurons accounted for the hyperactivity in the midbrain. Hierarchical clustering of neuronal activity with tail movements revealed distinct motor- and non-motor-associated hyperactive clusters in the hindbrain and midbrain, respectively. We identified the Mesencephalic Locomotor Region (MLR) sandwiched between these domains, with enhanced glutamatergic firing rate and cholinergic activation. Furthermore, we found that Telencephalic corticotropin-releasing factor b (crhb) expressing neurons play a crucial role in mediating stress-response to the tectum, which in turn triggers a cascade of neuronal hyperactivity downstream via MLR. These findings reveal a neural mechanism that links stress-induced sensory processing with motor control systems in the absence of regulatory feedback from catecholaminergic neurons, suggesting a direct, unmodulated pathway that bypasses typical inhibitory controls.

## Main

Dopamine (DA), noradrenaline (NA), and adrenaline, collectively referred to as catecholamines (CA), are synthesized via hydroxylation of tyrosine to L-dihydroxyphenylalanine (L-Dopa) by the tyrosine hydroxylase (TH) enzyme^1^ produced by catecholaminergic (CA) neurons. In mammals, these neurons form distinct populations in the midbrain (Ventral Tegmental Area - VTA^2^, Substantia Nigra - SNc^3^) and hindbrain (Locus Coeruleus - LC^4^). Loss of CA neurons has been linked to various neurodegenerative disorders, namely Parkinson’s (loss of SNc DA and LC NA neurons^5^) and Alzheimer’s (loss of VTA DA neurons^6^). In humans, about 60% Substantia Nigra (SNc) DA neurons are lost^7,8^ by the time motor symptoms necessitate PD diagnosis^9^. The motor symptoms include slowness of movement (bradykinesia), loss of movement (akinesia), rigidity, and postural instability, which collectively contribute to variability in gait and difficulty in walking^10^. Non-motor symptoms can precede the motor manifestations by several years and include olfactory loss (hyposmia), constipation, depression, and daytime sleepiness. However, even in advanced stages, PD patients can intermittently exhibit complex motor movements in response to an urgent stimulus^11–14^, including nightmares^15^. This phenomenon was termed paradoxical kinesia by Souques^16^ nearly a century ago, but remains poorly understood, despite some reports of compensatory mechanisms associated with PD or in animal models of PD^17,18^.

One of the earliest works^19^ reported paradoxical kinesia in dopamine-depleted rats, which, despite being akinetic in their home cages, swam effectively when placed in deep water or in a shallow floating ice bath. These behaviors were unaffected by dopamine antagonists, indicating that the paradoxical kinesia was not due to any residual dopamine. More recently, paradoxical kinesia was induced in haloperidol-treated rats using appetitive 50-kHz ultrasonic vocalizations and then disrupted by administering the glutamate agonist NMDA into the inferior colliculus^20^. This work suggests that the neural mechanisms involved in paradoxical kinesia are more glutamatergic than GABAergic.

Dopamine’s role in modulating the neural circuitry associated with locomotion, hunger, and motivation is evolutionarily conserved. In zebrafish, due to a genome duplication event at the base of teleost evolution, there exist two paralogous TH-encoding genes^21–24^: *TH1* & *TH2*. Mutants with developmental deficits of CA neurons^25,26^ display locomotion and reward-related behavioral deficits^27^. CA neuronal loss has also been induced post-development with neurotoxins or chemo-genetic ablation using the Nitroreductase (NTR)-Metronidazole (MTZ) system^28,29^, resulting in locomotive deficits, mimicking the phenotypes of PD^29–32^.

With an optically clear and millimeter-sized vertebrate brain, larval zebrafish are uniquely suited for whole-brain live imaging at cellular resolution^33^. Brain-wide calcium imaging with simultaneous recording of tail-swing behavior in head-constrained and tail-free larvae has provided important insights into the neural circuitry and its association with motor function^33–40^. Leveraging these advantages, we used transgenic zebrafish (nacre) larvae expressing TH1-Gal4 and TH2-Gal4 drivers along with UAS-NTR-mCherry and Elavl3:H2B-GCAMP6s transgenes. We investigated how the loss of both *TH1-* and *TH2-*expressing neurons affects free-swimming behavior, head-constrained tail-free behavior, and whole-brain neuronal activity. We found that larvae that exhibited severe locomotor deficit during free-swimming behavior showed no signs of locomotor deficit during head-constrained tail-free behavior. We analyzed neural activity (number of active neurons, their firing rates, and cumulative ΔF/F) across 2158 anatomically defined regions from Z-brain^41^ and mapZebrain^42^ atlases with HCR markers and transgene expressions. We found that elevated sensory activation in the tectum can be mediated to the motor regions in the hindbrain via Mesencephalic Locomotor Region (MLR), which may explain how urgent stimuli can trigger paradoxical kinesia in CA-deficient states.

These findings reveal a circuit-level functional rewiring that enables movement despite CA deficiency, with potential implications for understanding and treating motor symptoms in PD.

## Results

### Characterization and Ablation of CA Neurons

We generated zebrafish larvae expressing four transgenes: *TH1-gal4*^43^, *TH2-gal4-VP16*^32^, *UAS:NTR-mCherry*^28,44^ and *Elavl3:H2B-GCaMP6s*^45^ (**Fig. 1a**). At 4 days post-fertilization (dpf), larvae were sorted using stereomicroscopy and divided into two groups: control (0.1% DMSO) and ablated (5 mM MTZ with 0.1% DMSO). In the latter group, neurons expressing TH1-Gal4::UAS-NTR-mCherry or TH2-Gal4::UAS-NTR-mCherry are chemo-genetically ablated due to NTR-MTZ mediated cell-specific toxicity, which primarily acts through mitochondrial damage^31^. After this treatment, both groups were subjected to 24-48 hours of drug-free recovery in fish-system water, before being prepared for whole-brain two-photon imaging on 7 dpf to 8 dpf. Brainwide H2B-GCaMP6s signals (green channel) enabled alignment of mCherry expression (red channel) to a reference 3D stack (**Extended Data Fig. 1**) using ANTs^46^. We categorized TH1- and/or TH2-expressing neurons into 12 anatomical groups spanning 57 atlas-defined regions (**Supplementary Table 1**).

**Figure 1.**
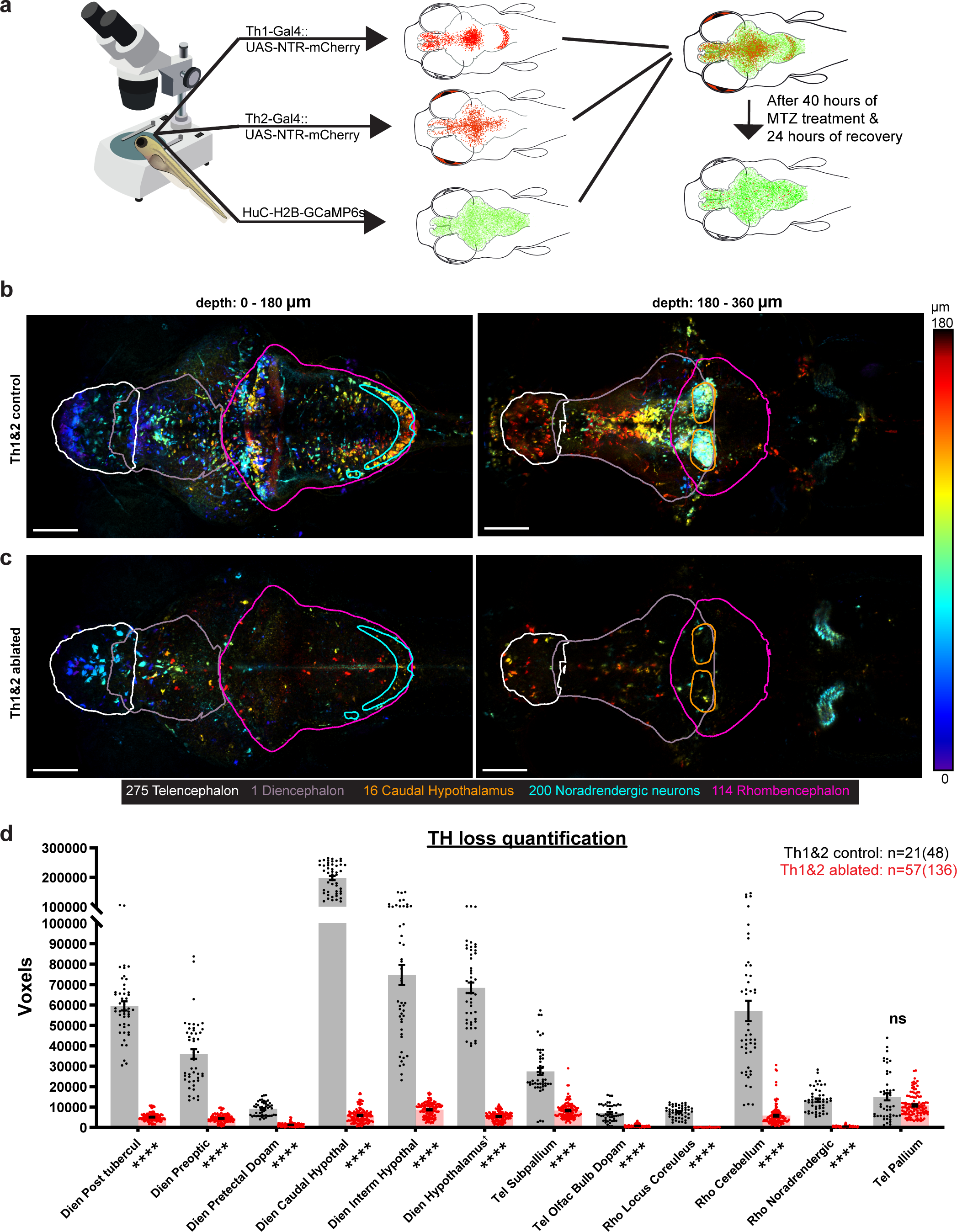
Chemogenetic ablation of catecholaminergic neurons using the NTR-MTZ system. **(a)** Schematic representation of transgenic lines showing expression patterns: TH1-Gal4::UAS-NTR-mCherry (top), TH2-Gal4::UAS-NTR-mCherry (middle), and pan-neuronal Elavl3:H2B-GCaMP6s (bottom). Catecholaminergic neurons expressing Th1 and Th2 were selectively ablated usin metronidazole (MTZ). **(b,c)** Maximum intensity projections of two-photon imaging data from control (b) and MTZ-treated (c) larvae following ANTs registration. Images are divided into dorsal and ventral halves and depth-coded for better visualization. Both Z-stacks were processed with the same intensity ranges before depth coding. Anatomical regions are delineated by contours: telencephalon (white), diencephalon (grey), rhombencephalon (pink), and Z-Brain-defined dopaminergic regions (red). Scale bar: 100 µm; field of view: 1 mm x 0.5 mm. Note: The posterior horn-like structure, located outside the brain, is more visible in (c) because higher PMT gains were typically used to visualize the red channel in the ablated groups. **(d)** Quantification of catecholaminergic neurons across 12 anatomical regions defined by the Z-Brain atlas. The bar graph shows mean voxel counts with individual data points overlaid (n = 48 Z-stacks from 21 control larvae; n = 136 Z-stacks from 57 MTZ-treated larvae). Asterisks adjacent to region-names bars indicate statistical significance (Wilcoxon rank-sum test and Permutation test: **\*\*\*\*** for p<0.0001 and **ns** for p≥0.05). †Caudal and Intermediate hypothalamus were removed from Hypothalamus.

In control larvae, mCherry expression driven by both Th1-Gal4 and Th2-Gal4 (**Fig. 1b, 1c**, **Supplementary Fig. 1**) was detected throughout the brain, with predominant expression in the hypothalamus. The caudal hypothalamus contained approximately three times more CA neurons than the intermediate hypothalamus, the next most CA-rich region. Significant populations of Th1-expressing neurons (**Supplementary Fig. 2**) were identified in the cerebellum, posterior tuberculum, noradrenergic region, locus coeruleus, subpallium, pretectum, and olfactory bulb (**Extended Data Fig. 2**). Additionally, a substantial population of Th2-expressing neurons was found in the Pallium and Preoptic area (**Fig. 1d**). Notably, we identified a previously unreported significant population of Th2-expressing neurons in the Tectum (**Supplementary Fig. 3**), which is not shown in **Fig. 1d**, as the zebrafish mesencephalon has been previously reported to be devoid of TH neurons in teleosts despite large populations present in the mammalian mesencephalon^47–49^ (particularly SN and VTA).

Two-photon imaging of larvae treated with MTZ revealed sparse populations of mCherry-labeled CA neurons, with many appearing structurally compromised (**Fig. 1c**, **Supplementary Fig. 4**, **Supplementary Fig. 5**). The loss was particularly evident in the dorsal-most noradrenergic neurons (z-brain id: 200) and the ventral-most caudal hypothalamic neurons (z-brain id: 16), which are delineated in cyan and orange, respectively, in **Fig. 1b and Fig. 1c**. These two regions also aided the verification of alignment accuracy in controls.

To quantify the loss of CA neurons due to chemogenetic ablation, voxel-based quantification was implemented, as cellular segmentation was particularly challenging in TH-dense regions. We counted the voxels above a certain threshold (iteratively determined) overlapping with 12 anatomical mask-composites (**Supplementary Table 1**) from 21 Th1&2 control (n=48 stacks), 57 Th1&2 MTZ-treated (n=136 stacks), five Th1-only control (n=10 stacks) and 16 Th1-only MTZ-treated (n=60 stacks). Statistical comparison of voxel counts using Wilcoxon rank-sum test and Permutation test demonstrated significant reduction (90-98%) in CA neurons across 11 of the 12 analyzed CA groups. Pallium was the sole region with non-significant loss of Th1- and Th2-expressing neurons. Interestingly, when we ablated only Th1-expressing neurons, the pallium had significant increase in voxel count (**Extended Data Fig. 2**).

### Locomotor deficits in freely swimming CA-deficient larvae

We investigated the behavioral consequences of CA neuronal ablation by analyzing free-swimming locomotion in 7-8 dpf larvae (**Fig. 2**). Using a custom-built setup containing 24 square wells (6" x 9" array, **Fig. 2a**), we recorded and tracked 870 larvae for 15 minutes each, following a 20-minute acclimation period. Larval tracking was accomplished using an in-house MATLAB program that automatically detected nacre larvae by their eyes, with a distinct contrast of at least 10 pixels against the white-light background. Following 40 hours of treatment with MTZ or DMSO (**Fig. 2b**), larvae were maintained in fish-system water for at least 24 hours before behavioral testing, which was conducted in fish-system water. Larvae with developmental defects or non-responsive to sudden tapping of the behavioral chamber were excluded from the behavioral assay. These larvae predominantly belonged to the ablated group. No feeding was done until all behavioral experiments were concluded.

**Figure 2.**
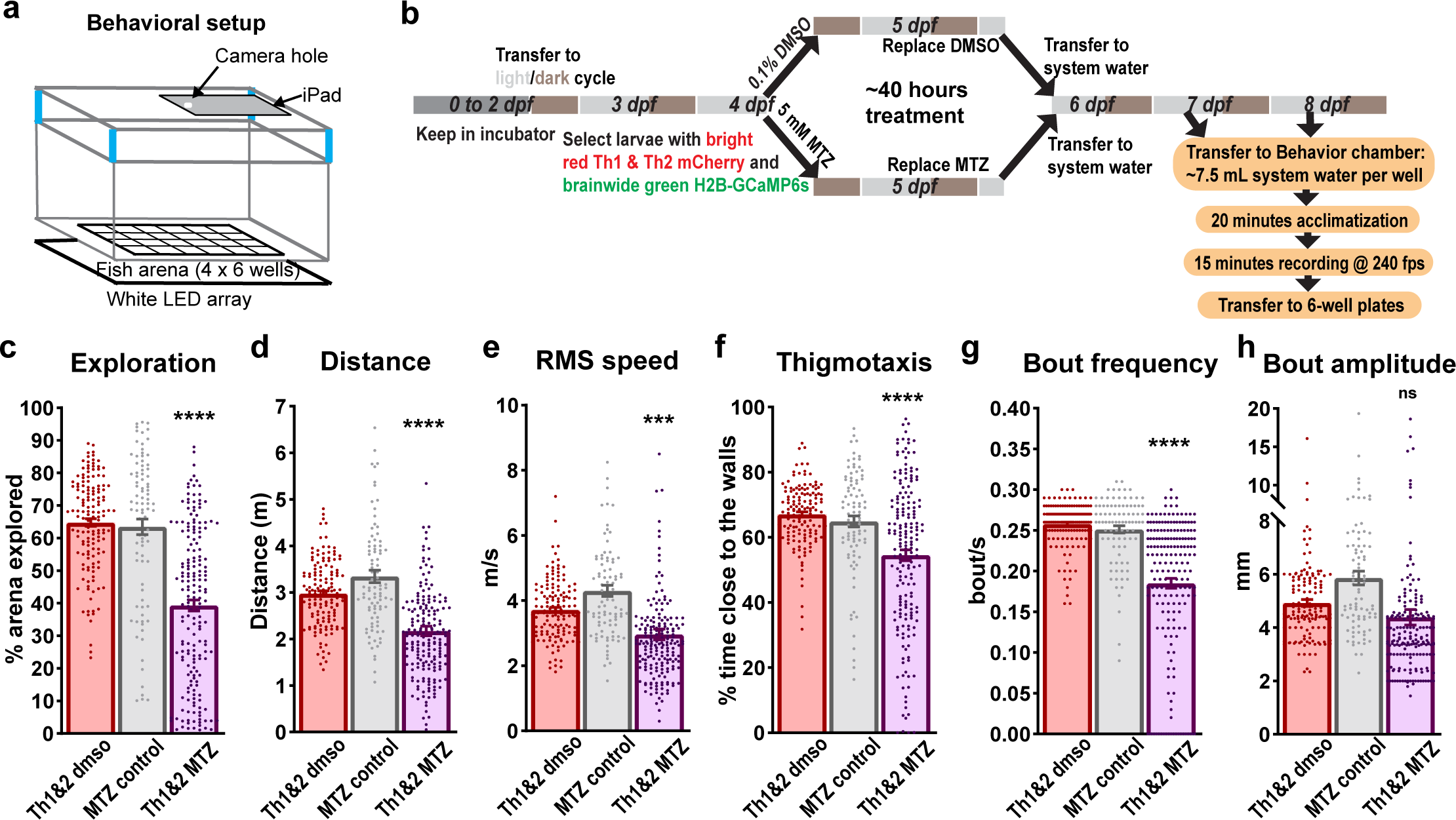
Free-swimming behavioral defects due to CA loss. **(a)** Behavioral setup used for recording free-swimming behavior of larvae, each with a 35 x 35 mm arena. **(b)** Flowchart depicting the experimental paradigm used for investigating the effect of CA loss (by NTR-MTZ mediated ablation) on free-swimming behavior of 7-8 dpf larvae. **(c)** Bar graph overlaid with individual datapoint showing the percentage of arena explored in 15 minutes by distinct groups of larvae: Th1&2 DMSO control (n=144, red), non-NTR MTZ control (n=90, grey), and Th1&2 MTZ treated (n=190, purple). “Th1&2” larvae were homozygous in three transgenes, viz., *Th1-Gal4*, *Th2-Gal4-VP16,* and *UAS-NTR-mCherry*. MTZ-treated larvae explored significantly less arena compared to their siblin controls treated with 0.1% DMSO and non-sibling MTZ controls, which did not express NTR-mCherry. Similar results were obtained for **(d)** total distance travelled in 15 minutes, **(e)** root-mean-square speed, **(f)** Thigmotaxis, and **(g)** frequency of bouts. **(h)** Interestingly, the distance covered in each bout (referred to as bout amplitude) was unaffected by the ablation of CA neurons. Th statistical significance was determined using ordinary one-way ANOVA followed by Tukey’s multiple comparison test, depicted a **\*\*\*\*** for p<0.0001, **\*\*\*** for p<0.001, and **ns** for p≥0.05 on top of the purple bar.

The study population comprised several experimental groups: 90 non-NTR-expressing controls treated with 5 mM MTZ, 138 larvae homozygous for Th1-Gal4 and UAS-NTR-mCherry, and 334 larvae homozygous for all three transgenes (Th1-Gal4, Th2-Gal4-VP16, and UAS-NTR-mCherry; represented in **Fig. 2c-h**). An additional 308 larvae, consisting of both homozygous and heterozygous carriers, underwent stringent fluorescence sorting at 4 dpf, with a 70-80% rejection rate for weaker Th1 and Th2 expression. The purely homozygous group was divided into control (n=144; 0.1% DMSO) and ablation groups (n=190; 5 mM MTZ in 0.1% DMSO), while the mixed carrier group was split into control (n=84; 0.1% DMSO), partial-ablation (n=65; 2.5 mM MTZ in 0.1% DMSO), and ablation (n=126; 5 mM MTZ in 0.1% DMSO) groups. These experiments spanned three years and multiple zebrafish generations, due to challenges of high mortality rates in Nace/Casper lines homozygous for multiple transgenes.

MTZ-treated Th1&2 larvae exhibited significant behavioral deficits compared to both DMSO-treated siblings and MTZ-treated non-transgenic controls. The CA-ablated larvae showed reduced arena exploration (**Fig. 2c**), decreased total distance traveled (**Fig. 2d**), lower root-mean-square speed (**Fig. 2e**), and reduced bout frequency (**Fig. 2g**). Notably, ablated larvae spent less time near the boundary walls (<6.75 mm from edges; **Fig. 2f**), suggesting potential alterations in anxiety-like behavior or spatial awareness. However, the distance covered per bout (bout amplitude) remained unchanged following MTZ treatment (**Fig. 2h**), consistent with previous findings by McPherson et al^32^, suggesting normal muscle function.

The behavioral deficits were specific to complete ablation of both Th1- and Th2-expressing neurons. Neither Th1-only ablation nor partial ablation of Th1 &Th2 using 2.5 mM MTZ produced significant locomotor deficit (**Extended Data Fig. 3**). The severity and range of the behavioral phenotype in Th1&2 ablated larvae were evident from spatial distribution analysis, where heatmaps of positional dwelling time revealed that Th1&2 ablated larvae spent prolonged periods in restricted areas compared to controls (**Extended Data Fig. 4**), consistent with their reduced exploratory behavior.

These behavioral deficits, combined with the confirmed ablation of CA neurons (**Fig. 1**), demonstrate that the loss of both Th1- and Th2-expressing neurons significantly impairs locomotor function in larval zebrafish.

### CA-deficient larvae with severely impaired free-swimming behavior exhibit paradoxically increased struggle behavior during functional brain imaging

Following the free-swimming behavioral recording, individual larvae (irrespective of their free-swimming behavior) were mounted upright in a way that they could freely move the tail below the swim bladder while the head was restrained for brain imaging (**Fig. 3a**). We recorded a total of 66 larvae (7-8 dpf) for a period of 24 minutes each: non-NTR MTZ control (n=7), Th1 control (n=2), Th1&2 control (n=13), Th1 treated with 5 mM MTZ (n=8), Th1&2 treated with 5 mM MTZ (n=30), Th1&2 treated with 2.5 mM MTZ (n=6).

**Figure 3.**
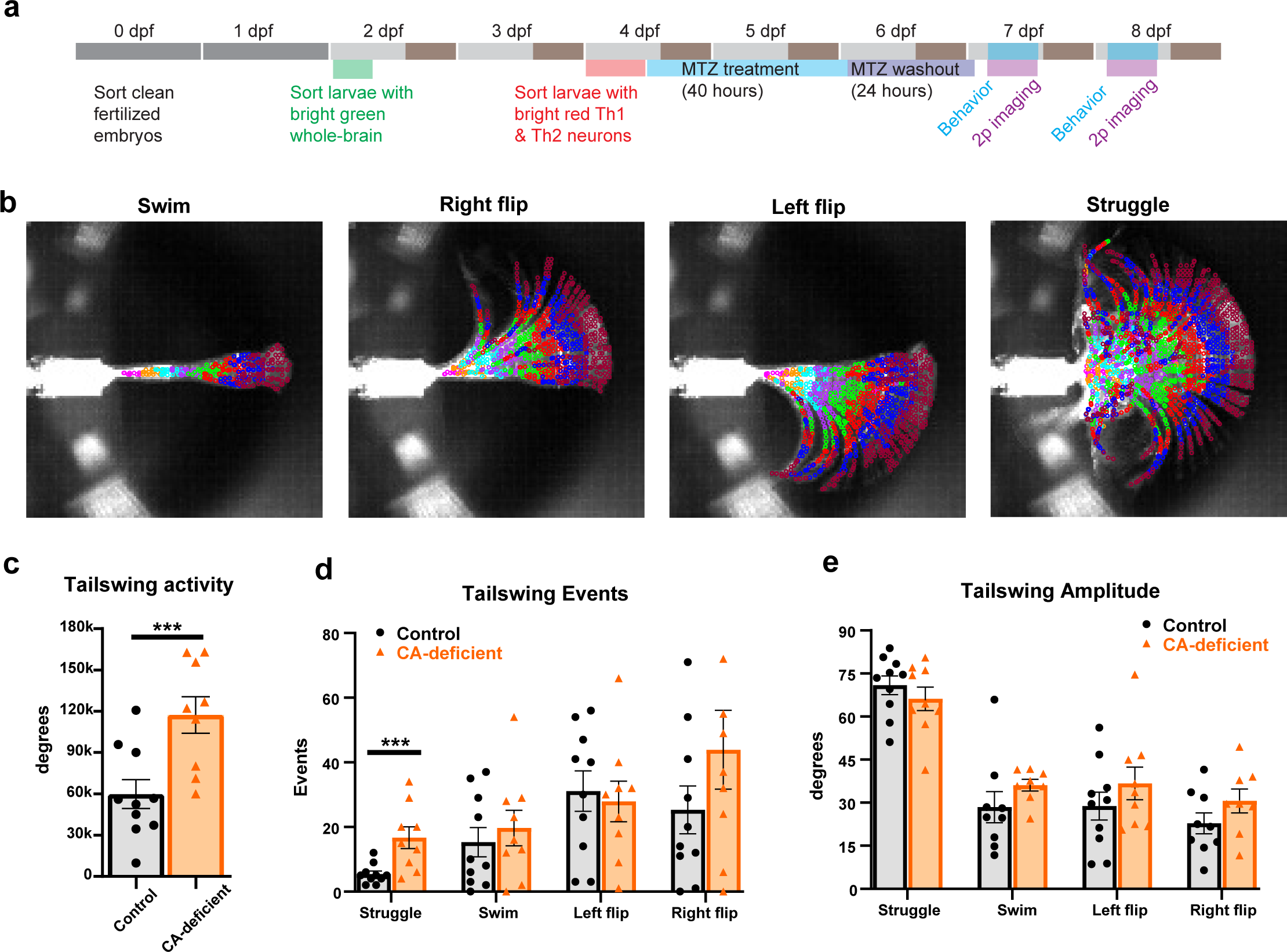
Paradoxical increase in tail movements of CA-deficient free-swimming impaired larvae during two-photon imaging. **(a)** Flowchart depicting the experimental paradigm used for investigating the effect of CA loss on head-constrained tail-free behavior of 7-8 dpf larvae combined with two-photon brain imaging. **(b)** Representative maximum intensity projections showing distinct tail movement patterns: swim, right-flip, left-flip, and struggle. Automated tail detection divided the tail into eight segments (color-coded: magenta, orange, cyan, purple, green, red, blue, and cherry). **(c-e)** Quantification of tail movements in control (n=10) and free-swimming CA-deficient (n=9) larvae during two-photon imaging: **(c)** frequency of tail-swing events, **(d)** total duration of each movement type, and **(e)** cumulative tail activity (sum of tail-swing angles) for struggle, swim, left flip, and right flip behaviors. Statistical significance determined by Wilcoxon rank-sum test (*** p<0.001).

We encountered significant heterogeneity in the free-swimming behavior of Th1&2 larvae treated with 5 mM MTZ, including those which were purely homozygous for both Th1 and Th2 expression (**Extended Data Fig. 4**). We focused on MTZ-treated larvae which exhibited significant free-swimming locomotor deficit, defined by a 35% cut-off for arena exploration in free-swimming behavior. 41% (n=130) of all Th1&2 larvae treated with 5 mM MTZ (n=316) were classified as “impaired” in free swimming, henceforth referred to as “CA-deficient” in this paper for simplicity.

To analyze tail movements during 2p-imaging, we developed a tail-tracking pipeline capable of detecting and skeletonizing the tail even when movements were faster than the imaging frame rate of ∼200 Hz and appeared blurry. We classified tail movements into swim, right flip, left flip, and struggle based on the average angle of segments four to seven (purple, green, red, blue; **Fig. 3b**).

Analysis of cumulative tail-swing angle (**Fig. 3c**) throughout the imaging period revealed that Th1&2 impaired larvae (n=9) showed 96% more tail activity compared to sibling controls (n=10; p=0.003). One control larva was excluded as an outlier (>3 scaled median absolute deviations from median; n=11 total), while no outliers were detected in the impaired group. Further analysis of tail-swing behavior showed that Th1&2 impaired larvae exhibited 209% more struggle events (p=0.0035) compared to controls, while forward swim, left flip, and right flip frequencies remained unchanged (**Fig. 3d**, and **Supplementary Fig. 6**). The average amplitude of tail swings was unaffected by CA loss across all movement types (**Fig. 3e**). Statistical significance was determined using Wilcoxon rank sum test (p<0.05).

These results suggest that CA-deficient zebrafish larvae with severe free-swimming locomotor deficit exhibit paradoxically increased tail-swing behavior (struggle) during two-photon brain imaging in head-restrained state.

### Brain-wide counts of active neurons, **Δ**F/F, and firing rates in free-swimming CA-deficient larvae compared to controls

To investigate the neural basis of elevated struggle behavior in free-swimming CA-deficient larvae during two-photon imaging, we analyzed whole-brain calcium activity in neurons expressing Elavl3-H2B-GCaMP6s (**Fig. 4a**). We employed suite2p (ver. 0.7.5) combined with custom MATLAB codes to extract neuronal positions and ΔF/F signals, remove false positives, and merge incorrectly split neurons. Neuronal spikes were inferred from Suite2p’s raw fluorescence traces using CNMFe’s constrained FOOPSI AR1 model (solved by CVX ver.2.1).

**Figure 4.**
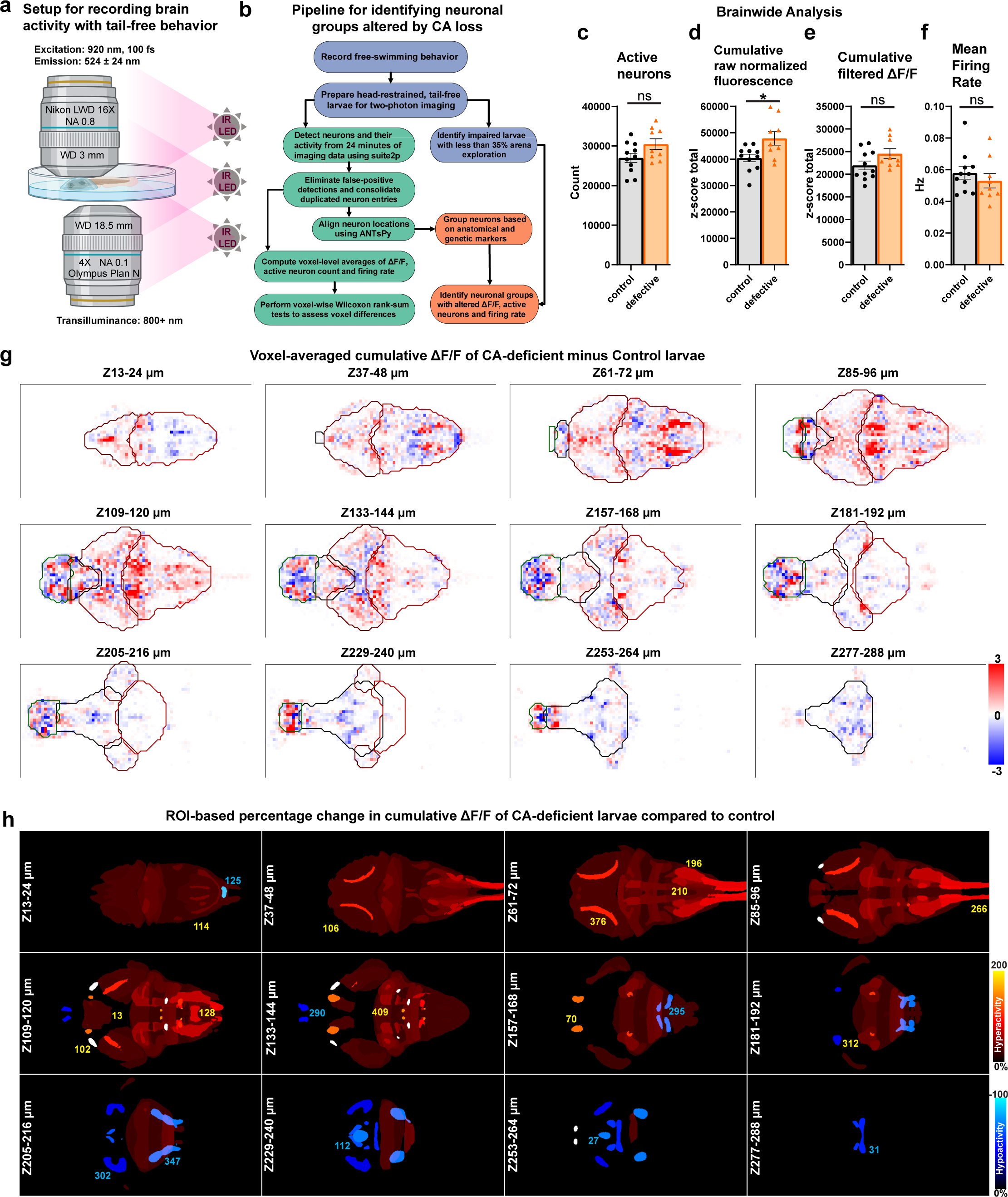
Altered Brain Activity in CA-deficient Larval Zebrafish. **(a)** Schematic representation of the imaging setup used to record neural activity simultaneously with tail movements in 7-8 days post-fertilization (dpf) zebrafish larvae, which were head-restrained using low-melting agarose after a free-swimming behavioral assay. **(b)** Flowchart illustrating the computational pipeline for estimating voxel-based and region-of-interest (ROI)-based neural activity differences between control and CA-deficient larvae from timelapse whole-brain calcium imaging data. **(c-f)** Bar graphs showing brain-wide neural activity metrics: **(c)** total number of active neurons, **(d)** total normalized fluorescence, **(e)** total cumulative ΔF/F, and **(f)** average firing rate. **(g)** Voxel-based comparison of z-scored cumulative ΔF/F revealed significant local variations between control and CA-deficient groups. Increased neural activity is indicated by red pixels, while reduced neural activity is indicated by blue pixels. The colorbar was saturated at ±3 to enhance contrast. **(h)** ROI-based view of anatomical regions showing altered cumulative ΔF/F in CA-deficient larvae compared to controls. The percentage increase is color-coded in warm colors (black-red-yellow-white), while percentage decrease is color-coded in cool colors (black-blue-cyan). Each panel in the 4 × 3 montage is a 12-µm maximum projection. Numeric labels shown in yellow and cyan fonts correspond to specific hyperactive and hypoactive regions, respectively. **(g,h)** Field-of-view: 0.5 mm (vertical) × 1.0 mm (horizontal).

The analysis revealed comparable numbers of active neurons between groups: 26,991 (±14.2%, n=11) in controls versus 30,460 (±13.4%, n=9) in free-swimming CA-deficient larvae (**Fig. 4b**). This difference was not statistically significant by either Wilcoxon rank-sum (p=0.111) or permutation tests (p=0.068). Similarly, the total z-scored cumulative ΔF/F showed no significant difference (p=0.097) between control (21,941 ±14.7%) and CA-deficient larvae (24,548 ±13.4%) (**Fig. 4c**).

Average firing rates per neuron (**Fig. 4f**) were calculated as the total number of peaks in the calcium signals divided by total recording duration (1445 seconds), yielding comparable (p=0.441) values: 0.058 Hz (±22.4%) for controls and 0.053 Hz (±25.7%) for free-swimming CA-deficient larvae. While suite2p’s native firing rate estimates were considerably higher (0.19 Hz for both groups), these were determined to contain numerous false positives and, thus, were not used.

The total z-scored cumulative raw fluorescence (0-1 normalized but without dynamic baseline removal) showed an 18% increase (p=0.0334) in free-swimming CA-deficient larvae (47,859 ±16%) compared to controls (40,441 ±11.6%). This finding was corroborated by suite2p’s native raw fluorescence measurements, which showed a 17% increase (p=0.023) in free-swimming CA-deficient larvae (38,222 ±12.5%) compared to controls (32,595 ±12.3%) (**Fig. 4d**). These results suggest elevated neural activity in free-swimming CA-deficient larvae at the raw signal level (after 0-1 normalization), prompting us to investigate local neural activity patterns.

### Voxel-based analysis reveals large-scale alterations to neural activity in CA-deficient brains

To systematically analyze region-specific differences in neuronal activity, we performed whole-brain comparative analysis between free-swimming CA-deficient (n=9) and control (n=11) zebrafish larvae. Following alignment to our reference template, we generated downsampled 3D-heatmaps for the number of active neurons (**Extended Data Fig. 5a**), their z-scored cumulative ΔF/F (**Fig. 4g**), and average firing rates (**Extended Data Fig. 6a**). The heatmaps were averaged across all larvae, followed by estimation of the difference between control and free-swimming CA-deficient brains. Prior to estimation of the average heatmap difference, Wilcoxon rank-sum test (p<0.05) was applied to all voxels to find those which were significantly altered between control and CA-deficient larval groups. The 3D-heatmaps were then mapped to 704 predefined anatomical regions (294 Z-brain atlas regions, 114 mapZebrain atlas regions and 296 HCR markers/transgenic expressions) to estimate the percentage of significantly altered voxels in established brain regions (**Supplementary Table 3)** corresponding to the heatmaps of cumulative ΔF/F (columns C, E), number of active neurons (columns G, I) and firing rate (columns K, M). The percentage of altered voxels did not reflect the magnitude of the difference between control and CA-deficient larvae; hence, we summed up the corresponding heatmap difference for the significantly altered voxels. The absolute sum of all these six scores, both hyperactive and hypoactive, was used as a comprehensive “Relevance” score to sort the brain regions/markers affected by CA loss.

Analysis of broad neuroanatomical domains revealed distinct patterns of altered activity. GABAergic neurons showed the highest relevance score (2883), comprised of 5% voxels with increased and 1.3% voxels with decreased cumulative ΔF/F. This was followed by the rhombencephalon (5.3% increased, 1.5% decreased ΔF/F, score 1915), glutamatergic neurons (7.1% increased, 1.9% decreased ΔF/F; score 1871), mesencephalon (6.4% increased, 1.9% decreased ΔF/F; score 1396) and cholinergic neurons (8.3% increased, 1.6% decreased ΔF/F, score (1077). Diencephalon and Telencephalon were ranked 19^th^ and 20^th^ respectively in terms of their relevance scores with more balanced changes to cumulative ΔF/F.

At a finer anatomical scale, the tectum, pallium, cerebellum, and subpallium emerged as key regions of interest. The tectum and cerebellum showed pronounced hyperactivity (8.1% and 6.3% increased ΔF/F, respectively) with minimal decreases (1.5% and 1.0%). The pallium and subpallium displayed more complex patterns, with substantial bidirectional changes (pallium: 6.5% increased, 4.6% decreased; subpallium: 5.4% increased, 5.3% decreased). Of particular interest, crhb-expressing neurons showed marked hyperactivity (9.4% increased, 1.1% decreased ΔF/F).

These findings indicate predominantly heightened neuronal activity in the midbrain and hindbrain of free-swimming CA-deficient larvae. While most of the hyperactivity conformed to anatomically defined regions, hypoactive clusters were sparsely distributed particularly in the forebrain, but frequently with hyperactive clusters in proximity. While this downsampled analysis effectively revealed distinct regional activity patterns, the reduced spatial resolution limited detection of changes in smaller anatomical structures, prompting more precise analysis by directly labelling individual neurons (**Figure 4h**). However, this method provides a unique advantage of identifying novel brain regions altered by CA loss, which allowed us to identify the Mesencephalic Locomotor Region (MLR) (**Extended Data Fig. 7**), closely matched with the findings of Martin et al^50^. The complex local variations in voxel-based neural activity differences between CA-deficient and control larvae prompted us to compartmentalize the brain into 2158 regions derived from 704 predefined anatomical regions and markers from z-brain and mapZebrain.

### ROI-based analysis reveals that neuronal hyperactivity is accompanied by paradoxical reduction in firing rate

By registering the neuronal centroids to z-brain and mapZebrain, we identified 2158 neuronal groups, followed by estimation of their cumulative ΔF/F, number of active neurons, and average firing rate in CA-deficient (n=9) and control (n=11) zebrafish larvae. We found 204 neuronal groups with increased and 7 neuronal groups with decreased cumulative ΔF/F in the hindbrain (**Fig. 5a**, **Supporting Table 4**). In the midbrain, 77 neuronal groups had increased, and 5 neuronal groups had decreased cumulative ΔF/F (**Supporting Table 5**). The forebrain, however, showed significant complexity with increased ΔF/F in 23 neuronal groups and reduced ΔF/F in 30 neuronal groups (**Supporting Table 5**). Major hyperactive regions were Rhombencephalon (in hindbrain), Tectum (in midbrain) and dorsal thalamus (in forebrain). While the entire hindbrain of CA-deficient brains was mostly hyperactive, abducens motor nucleus and octaval ganglion had significantly reduced ΔF/F compared to healthy controls. Other hypoactive regions were far upstream of the hindbrain, primarily in the sub-pallium and hypothalamus.

**Figure 5.**
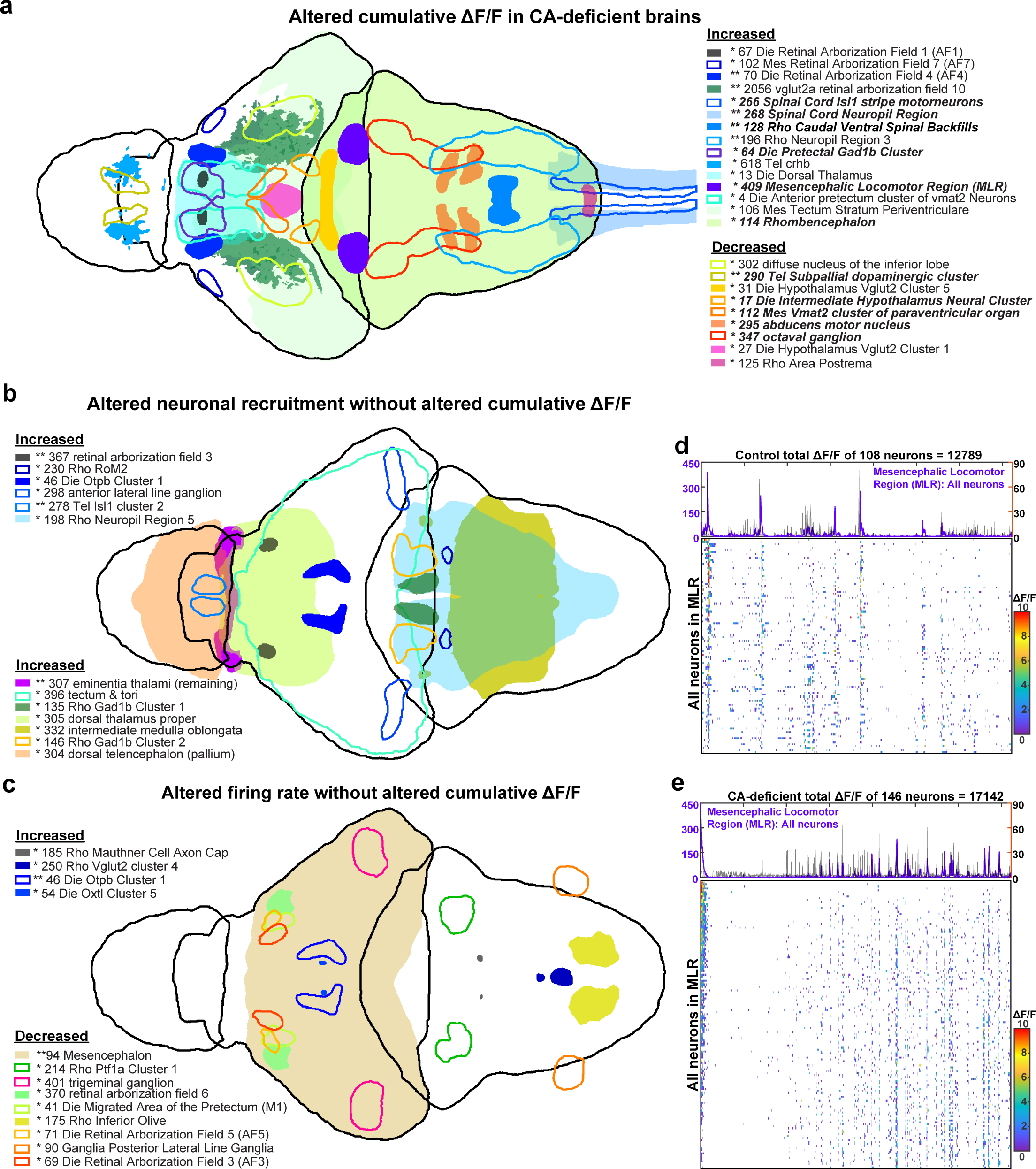
Brainwide hyperactivity in CA-deficient larvae is accompanied by paradoxical reduction in firing rate. **(a)** Primary brain regions with significantly altered cumulative ΔF/F in CA-deficient larvae. Many of these regions also had an increased number of active neurons and/or altered firing rates. Regions with reduced cumulative ΔF/F are shown in warmer colors. Regions in bold italic font were found to be associated with tail movements. **(b)** Primary brain regions which had increased number of active neurons without significant increase in cumulative ΔF/F. No such regions were identified with reduced number of active neurons. Many of these regions had reduced firing rates. **(c)** Primary brain regions with reduced firing rate and unaltered cumulative ΔF/F. Some of these regions had altered number of active neurons. In panels a-c, sub-regions with similar hyperactivity or reduction in firing rate have not been shown to avoid crowding. **(d,e)** Temporal ΔF/F traces of all neurons in the Mesencephalic locomotor region (MLR) of a representative **(d)** control and **(e)** CA-deficient brain. The summed ΔF/F is shown on top in purple color (same as corresponding mask color in panel **a**) and overlaid with tail-angle timeseries (black trace, range 0° to 90°) represented by the absolute angle (degrees) of the tail from the dynamically estimated resting position. The horizontal axis represents time (also for raster plots). The title of each plot shows the number of active neurons and their normalized cumulative total ΔF/F across the entire recording duration (1445 seconds). The colormap used for the raster plots have the range of 0-10 as depicted in the colorbar, where the minimum (0) is represented as white background, while ΔF/F values ≥10 are represented by red color. Both raster plots have the same vertical resolution (2 units per neuron), so the height of the raster plots is proportional to the number of active neurons. Plotted ΔF/F values were z-score normalized followed by subtraction with minimum to make all values positive.

Increase in cumulative ΔF/F was found to be frequently associated with increased neuron activation in the corresponding hyperactive neuronal group. We found that 101 out of the 204 hyperactive neuronal groups in the hindbrain had significantly more active neurons in CA-deficient larvae compared to control. Surprisingly, none of the 204 hyperactive neuronal groups had increased firing rates, while four had reduced firing rates (p<0.05). Similarly, none of the 77 hyperactive neuronal groups in the midbrain had increased firing rate, while 33 hyperactive neuronal groups had significantly reduced firing rate.

Increased neuron activation was not always sufficient to significantly increase cumulative ΔF/F as observed in 52, 29, and 35 neuronal groups across the hindbrain, midbrain, and forebrain, respectively, with a significant increase in the number of active neurons but unaltered cumulative ΔF/F (**Fig. 5b**). Pallium was among the largest structures identified with this characteristic. This might be due to a decrease in firing rates, which was statistically significant across 14 neuronal groups in the midbrain and one in the forebrain (**Fig. 5c**), but with unaltered cumulative ΔF/F despite a significant increase in the number of active neurons. Increase in the firing rate alone was never found to be sufficient to increase the cumulative ΔF/F of any neuronal group.

### Free Swimming-CA-deficient larvae show distinct neurotransmitter-specific alterations of activity

GABAergic neurons (gad1b), which accounted for 17.6% of total hindbrain volume, had 31% higher (p=0.028) cumulative ΔF/F. Cholinergic (chata) neurons were the second largest neuronal group in the hindbrain (6.6% by volume) with 41% higher (p=0.002) cumulative ΔF/F and 16% higher (p=0.048) number of active neurons. The glutamatergic domain in the hindbrain (3.2% by volume) had 46% higher (p=0.015) cumulative ΔF/F and 23% higher number of active neurons (p=0.028). Among other notable hyperactive neuronal groups in the hindbrain were otpa, otpb, gata2a, vagus motor nucleus, spinal cord Isl1 stripe motoneurons, and mesencephalic locomotor region (**Fig. 5a**).

In the midbrain of CA-deficient larvae, primarily in the tectum, GABAergic neurons had 27% higher cumulative ΔF/F (supplementary table 4), 23% higher number of active neurons, and 28% lower firing rate. Glutamatergic neurons were the second largest group of hyperactive midbrain neurons, had 31% higher cumulative ΔF/F, 33% higher number of active neurons and 24% lower firing rate. Midbrain cholinergic neurons were not hyperactive despite 18% higher number of active neurons, albeit with 23% lower firing rate.

In the forebrain, the three major neurotransmitters overall had unaltered cumulative ΔF/F, number of active neurons, or firing rate, but had complex local variations. For instance, glutamatergic neurons in the dorsal thalamus had a 58% higher cumulative ΔF/F and 35% higher number of active neurons, while major pockets of hypothalamic glutamatergic neurons had significantly reduced ΔF/F. We found that forebrain corticotropin-releasing factor b (crhb) expressing neurons, co-expressing vglut2a, had 57% higher cumulative ΔF/F (p=0.005), 32% higher active neuron count (p=0.003), and 31% lower firing rate (p=0.019). Hyperactive GABAergic neurons in the forebrain were primarily localized in the Pretectal Gad1b cluster and the Anterior pretectum cluster of vmat2 neurons. On the other hand, GABAergic neurons in the subpallial dopaminergic cluster and Olig2 cluster had significantly reduced cumulative ΔF/F.

### Most of the Retinal Arborization fields are Hyperactive in CA-deficient Larvae

The retinal arborization fields in the midbrain (**Supplementary Table 5**) and forebrain (**Supplementary Table 6**) were found to be significantly more active in CA-deficient larvae compared to those in control brains. Retinal AF10 was the largest arborization field, which, despite not being significantly altered overall, had glutamatergic neurons (**Fig. 5a**) with 159% higher cumulative ΔF/F (p=0.008), 109% increased number of active neurons (p=0.008), and 33% lower firing rate (p=0.01). *Retinal AF7* was the second largest midbrain arborization field, 2.9 times smaller than vglut2a Retinal AF10, with 163% higher active neurons (p=0.0043) and 311% higher cumulative ΔF/F (p=0.0135).

In the forebrain, *Retinal AF4* had 172% higher (p=0.006) cumulative ΔF/F accompanied by 70% increase in active neurons and 36% decrease in firing rate. Control larvae rarely had any neurons detectable in Retinal AF1 (0.5±2.3), closely followed by Retinal AF3 (0.8±3.2). In CA-deficient larvae, the number of active neurons rose by 413% and 294% in AF1 and AF3, respectively. This led to a 1300% increase in Retinal AF1’s cumulative ΔF/F. In AF3, the firing rate was reduced by 76% (p=0.004), which resulted in unaltered cumulative ΔF/F.

### Motor-associated hyperactive neuronal clusters are primarily localized in the hindbrain

To uncover the neuronal circuitry associated with tail movements, we employed hierarchical clustering based on Spearman correlation. The total number of active neurons associated with tail movements was 20255±4495 in control and 20768±3073 in CA-deficient brains. Analyzing the cumulative ΔF/F of these neurons across the 2158 brain regions/markers, we found elevated motor-associated neural activity in 147 hindbrain-neuronal groups (**Supplementary Table 7**), 17 midbrain-neuronal groups (**Supplementary Table 8**) and 8 forebrain-neuronal groups (**Supplementary Table 9**). On the other hand, 4, 4, 26 hypoactive motor-associated neuronal groups were identified in the hindbrain, midbrain, and forebrain, respectively. Most of the motor-associated hindbrain regions shown in **Fig. 6a** were identified earlier with altered cumulative ΔF/F (**Fig. 5a**, bold font).

**Figure 6.**
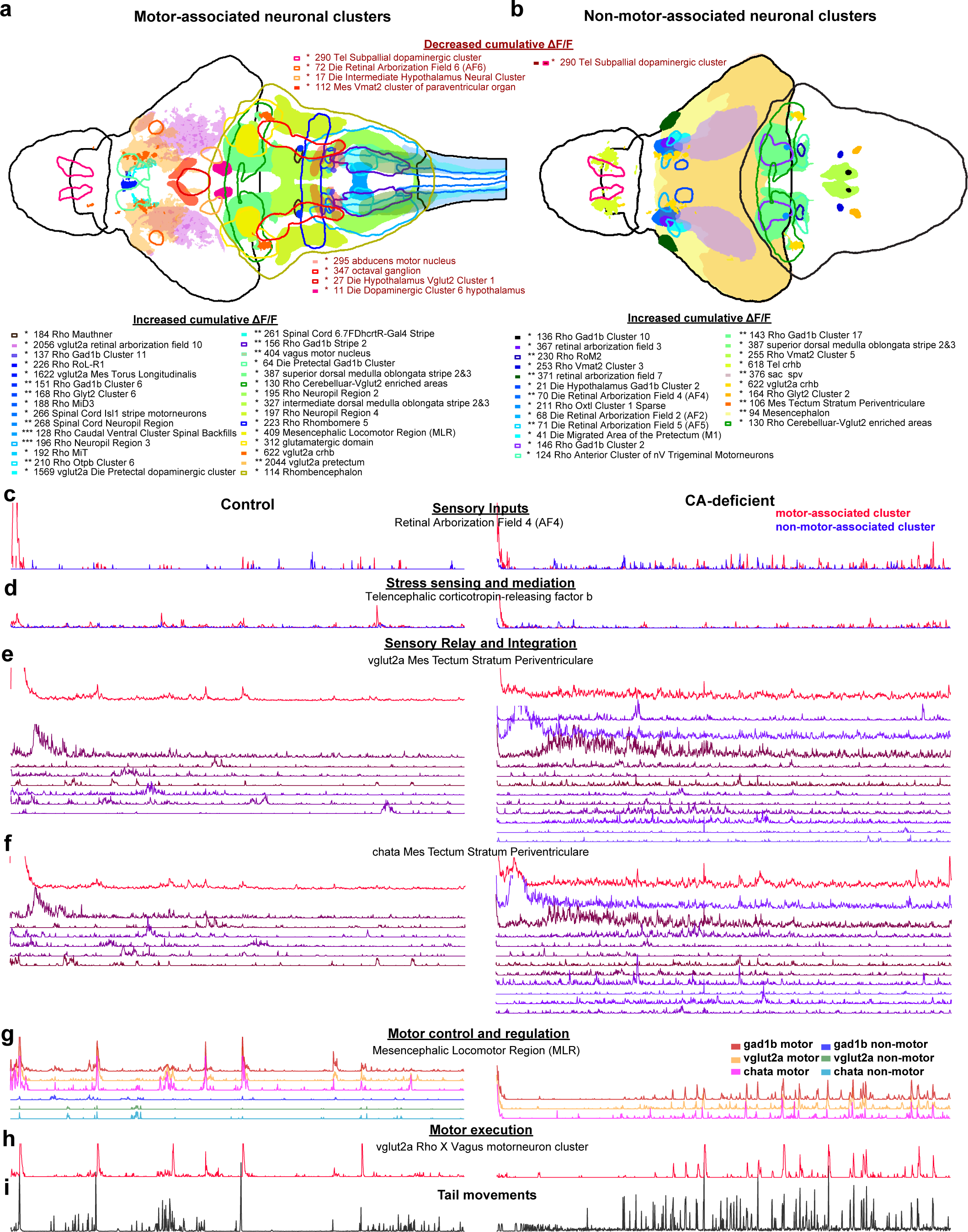
Motor-associated and non-motor-associated neuronal clusters have distinctive characteristics and distribution. **(a)** Brain regions with motor-associated neuronal clusters and **(b)** non-motor-associated neuronal clusters with increased or decreased (red font) cumulative ΔF/F in CA-deficient brains compared to control. Each region is represented by a unique color and either delineated or filled to enhance visibility, with corresponding region-names shown below. Not all regions have been shown to avoid crowding. **(c-h)** Temporal ΔF/F traces of a representative neuronal group from a control (left) and CA-deficient (right) larva. Red traces represent the summed ΔF/F of all motor-associated neurons in the corresponding region, while blue traces represent the summed ΔF/F of all remaining neurons in that region. The activity sum was divided by the total number of neurons in the region. The panels are arranged from top to bottom based on their hypothesized placement in the neural circuitry from **(c)** perception of stimuli to **(h)** execution of struggle behavior, all of which were hyperactive in CA-deficient brains. **(e,f)** Temporal ΔF/F traces (purple shades) of non-motor-associated neuronal clusters in the tectum (glutamatergic/cholinergic) have been separately shown based on their hierarchical cluster. **(g)** Temporal ΔF/F traces of motor-associated (red, yellow, magenta) and non-motor-associated (blue, green, cyan) neuronal clusters in the MLR have been shown separately based on their GABAergic, glutamatergic, and cholinergic labels. The specific CA-deficient larva shown here did not have non-motor-associated neurons in the MLR. **(i)** Tail movements of the corresponding control (left) and CA-deficient (right) larva shown as black trace, with vertical axis representing the absolute angular deviation from resting position. The horizontal axis represents time for all the traces (c-i) and are aligned.

In the midbrain, notable motor-associated hyperactive neuronal groups were primarily glutamatergic: vglut2a Die Pretectal dopaminergic cluster, vglut2a Mes Tecum Neuropil, vglut2a Mes Torus Longitudinalis, vglut2a vglut2a pretectum, vglut2a retinal arborization field 10 and vglut2a sac-spv. Hypoactive GABAergic neurons were identified in gad1b stratum marginale, while hypoactive cholinergic neurons were identified in chata stratum griseum centrale.

In the forebrain, hyperactive motor-associated neuronal groups were identified in the Pretectal Gad1b cluster, vglut2a crhb and vglut2a Preoptic area as regions. Motor-associated forebrain regions were primarily hypoactive, notably, the Intermediate Hypothalamus, the Mesencephalic Vmat2 cluster of paraventricular organ, the Subpallial dopaminergic cluster, and glutamatergic neurons in the Tel Anterior Commissure.

### Hyperactive neuronal clusters not directly associated with tail movements are primarily localized in the midbrain

Following the identification of neurons associated with tail movements, the remaining 25% neurons in control and 32% neurons in CA-deficient larvae were classified as non-motor associated neurons. Analyzing the cumulative ΔF/F of these neurons across the 2158 brain regions/markers, we found 75 hyperactive hindbrain-neuronal groups (**Supplementary Table 10**), 123 hyperactive midbrain-neuronal groups (**Supplementary Table 11**) and 32 hyperactive forebrain-neuronal groups (**Supplementary Table 12**). Additionally, six non-motor-associated neuronal groups were identified in the forebrain and one in the hindbrain, with reduced cumulative ΔF/F.

In the hindbrain, notable non-motor-associated hyperactive neuronal groups were Rho Cerebelluar-Vglut2 enriched areas (#130) and superior dorsal medulla oblongata stripe 2&3 (#387), both of which also had motor-associated neuronal groups with elevated cumulative ΔF/F. Hypoactive non-motor-associated GABAergic (gad1b) neurons were identified in gad1b Rhombomere 1 (#1786), gad1b Rhombomere 2 (#1789) and gad1b Rho Otpb Cluster 6 (#1771). Rho Vmat2 Cluster 5 and Rho Vmat2 Cluster 3 and Rho RoM2 were among other notable non-motor-associated hyperactive regions. Vglut2a olig3 was the only hypoactive non-motor-associated hindbrain neuronal group. Cerebelluar-Vglut2 enriched areas and superior dorsal medulla oblongata stripe 2&3 were found to be hyperactive for both motor-associated and non-motor-associated neuronal clusters.

The entire midbrain was found to be non-motor-associated with increased cumulative ΔF/F (**Fig. 6b**), comprised of GABAergic (#1591, p=0.0078), glutamatergic (#1590, p=0.005) and cholinergic (#1592, p= 0.0098) neurons. Sub-regions of interest include tectum, retinal arborization field 10, stratum fibrosum et griseum superficiale, Mes Medial Tectal Band, retinal arborization field 7, gad1b pretectum and vglut2a pretectum.

In the forebrain, Tel crhb (#618), vglut2a Tel Vglut2 rind (#1898), Retinal AF4, Die Migrated Area of the Pretectum (M1), Retinal AF2, Die Hypothalamus Gad1b Cluster 2 and gad1b Tel Anterior Commissure (#1876) were among the notable non-motor-associated hyperactive neuronal groups. Subpallial dopaminergic cluster in the telencephalon was found to be hypoactive for both motor-associated and non-motor-associated neuronal clusters.

### MLR sits at the crux of motor and non-motor associated neuronal clusters with enhanced glutamatergic and cholinergic signaling

To investigate the functional difference between midbrain and hindbrain with respect to association with tail movements, we aimed to identify the most upstream hyperactive hindbrain neuronal groups. This led to the identification of the glutamatergic domain (#312) and Mesencephalic Locomotor Region (#409), as shown **Fig. 6a**, sandwiched between the midbrain and hindbrain. We verified MLR’s association to tail movements by plotting temporal traces (ΔF/F) of all neurons overlaid on the tail-angle timeseries for control (**Fig. 5d**) and CA-deficient larvae (**Fig. 5e**).

Both the glutamatergic domain and Mesencephalic Locomotor Region (MLR) were found to be comprised of GABAergic, glutamatergic, and cholinergic neurons (**Supplementary Table 2**). The glutamatergic domain had increased cumulative ΔF/F for all as well as motor-associated neurons corresponding to vglut2a glutamatergic domain (#1934), gad1b glutamatergic domain (#1935) and chata glutamatergic domain (#1936). vglut2a glutamatergic domain also had an increased number of active neurons.

26% of MLR (10133 voxels) was found to overlap with the glutamatergic domain, but it was functionally different. Vglut2a MLR (#411) was found to have 44% higher firing rate (p=0.0062), while chata MLR (#416) was found to have 51% higher active neurons (p=0.0059). Neither vglut2a MLR nor chata MLR was significantly altered when motor-associated neurons were analyzed, while the entire MLR had 45% higher (p=0.048) ΔF/F corresponding to motor-associated neurons (**Supplementary Table 7**). GABAergic neurons in the MLR were very few, and with no significant change in any of the nine parameters analyzed (number of active neurons, cumulative ΔF/F, firing rate, cumulative ΔF/F only during tail movements, motor-associated ΔF/F, non-motor-associated ΔF/F, number of motor-associated neurons, fraction of motor-associated neurons and average correlation of motor-associated neurons with tail-swing angle).

The MLR is thus more complex compared to glutamatergic domain with unique neurotransmitter roles in compensating for CA loss (**Fig. 6g**). Additionally, by measuring the correlation between MLR neurons with those in other 2157 neuronal groups (**Supplementary Table 13**), we found that it has the highest connectivity with Otpb Cluster 2 locus coeruleus (#206), followed by RoM1 (#229). We did not detect any functional connection between MLR and Nucleus of the Medial Longitudinal Fasciculus (NMLF), verifying Martin et. al.’s findings^50^.

### The tectum and telencephalic crhb share a non-tail cluster of neurons that fire concomitantly following the onset of brain imaging

Both the tectum and Telencephalic crhb (corticotropin-releasing hormone b) were identified as hyperactive non-motor-associated regions (**Supplementary Table 11, 12**). Corticotropin-releasing hormone is known to play a crucial role in stress regulation^51^. The non-motor associated neurons in the tectum were found to form over a dozen distinct hierarchical clusters, some containing hundreds of neurons. We found that one of these clusters had an activity peak (hump) that temporally matched a similar peak in one of the non-motor-associated neuronal clusters in Tel crhb, within the first hundred seconds of two-photon imaging (**Extended Data Fig. 8**). This suggests that CA neurons have negative feedback on crhb expressing neurons of healthy controls.

By measuring the correlation between Tel crhb neurons with those in other 2157 neuronal groups, we found that it is connected to 86 brain regions, notably: retinal AF8, vglut2a dorsal thalamus proper, ventral thalamus, hypothalamus, pallium, subpallium, midbrain (particularly tectum) and the hindbrain (**Supplementary Table 14**). The connectivity with chata Mes Tectum Stratum Periventriculare (#1615) was severely weakened (-94%, p=0.0424).

These findings suggest that Telencephalic corticotropin-releasing factor b (crhb) expressing neurons play a crucial role in mediating stress-response to the tectum, which in turn triggers a cascade of neuronal hyperactivity downstream.

## Discussion

Chemogenetic ablation of both TH1 and TH2 expressing neurons, but not TH1-expressing neurons alone, induced free-swimming locomotor deficits in larval zebrafish, particularly reduced arena exploration. Paradoxically, when these CA-deficient larvae were imaged under a two-photon microscope in a head-restrained tail-free preparation, their cumulative tailswing angle doubled compared to control siblings. We identified hyperactive motor-associated neuronal clusters predominantly in the hindbrain and hyperactive non-motor associated neuronal clusters localized in the midbrain and forebrain. The Mesencephalic Locomotor Region (MLR) emerged as a key motor-associated region, showing enhanced glutamatergic firing rate and increased activation of cholinergic neurons, resulting in elevated cumulative ΔF/F.

In both control and CA-deficient larvae, the MLR is functionally connected to the tectum, a key non-motor-associated hyperactive region in CA-deficient larvae with reduced firing rate. The tectum was found to be functionally connected to corticotropin-releasing factor b (crhb) expressing neurons in the Telencephalon (Tel crhb), which in turn has strong connectivity with the glutamatergic domain adjacent to MLR with significant overlap. Although direct connectivity between MLR and Tel crhb was not observed, these regions share 33 common connections, with the glutamatergic domain being the only sub-region showing hyperactive motor-associated neurons in CA-deficient larvae. This led to the identification of a feedback circuit entangling motor-associated neurons in the hindbrain with non-motor-associated neurons in the midbrain and forebrain.

During imaging, larvae experience vibrational disturbances from objective lens movement and focused photothermal irritation from the femtosecond laser beam. The zebrafish larva is also basked in infrared light used for motion tracking, which is visible to the human eyes as a pinkish hue. The multimodal sensory integration of these stimuli in the tectum via hyperactivation of retinal arborization fields in the absence of catecholaminergic (CA) neurons is the primary driver for paradoxical kinesia. The presence of intense multimodal (vibration, heat, infrared light) sensory stimuli can bypass crhb-expressing neurons and lead to vigorous swinging of the tail at the onset of two-photon imaging^52^. It is so commonly observed that whenever we do not see it, we suspect that larvae have gotten injured during the mounting procedure, and brain imaging is not pursued further. While crhb-expressing neurons show hyperactivity in CA-deficient larvae, this likely indicates unregulated stress levels rather than directly driving paradoxical kinesia. Individual variation in coping responses to stress with relation to dopamine is very well documented^53–55^.

The mesencephalic locomotor region (MLR) is a conserved brainstem hub across species, involved in controlling locomotion and integrating behavioral and autonomic responses. In lampreys, zebrafish, and amphibians^56^, the MLR reliably triggers rhythmic movement, reflecting its evolutionary origin. Rodent studies using modern tools like optogenetics show that the cuneiform nucleus (CnF) drives fast locomotion, while the pedunculopontine nucleus (PPN) modulates slow movement and behaviors like arousal and grooming^57^. In cats and primates, MLR stimulation influences gait and posture, while in humans, clinical trials using deep brain stimulation (DBS)^58,59^ target the MLR (mostly PPN or CnF) to treat Parkinsonian gait disturbances^60,61^, though with variable success. Based on our findings and reported clinical trials, we propose that the overlapping region between the MLR and Glutamatergic domain could be a more promising target for stimulation than the currently targeted PPN (in MLR) to alleviate motor deficits induced by the loss of CA neurons.

We developed several technical innovations to enable these discoveries. These include high-throughput behavioral tracking optimized for transparent larvae, efficient tail movement analysis at lower frame rates, and improved anatomical registration that combines multiple zebrafish brain atlases. Our integration of suite2p and CNMF for calcium imaging analysis with an additional layer to remove false positives and merge wrongly split neurons, provides enhanced accuracy in neuron detection and signal extraction. These tools collectively enabled the largest ever reported brainwide functional classification of the zebrafish brain and are now available to the entire research community to enable more precise investigation of neural circuits across model organisms.

## Code and Data Availability

All MATLAB, python and ImageJ-macro scripts developed for this project can be downloaded from https://github.com/DrKrishB/THlossCompensation. It also contains some data and results reported in this paper as examples.

## Methods

### Zebrafish lines and screening

All procedures were approved by the University of California San Francisco Institutional Animal Care and Use Committee (IACUC). Zebrafish embryos were maintained in E3 medium (5 mM NaCl, 0.17 mM KCl, 0.33DmM CaCl_2_, 0.33DmM MgSO_4_) till 6 dpf and then in fish-system water. Embryos were incubated at 28^°^C in dark chamber for 0-2 dpf and then transferred to fish room maintained at 27°C and 14/10-hour light-dark cycle.

Four transgenic lines from four different labs were combined into a single transgenic nacre zebrafish line carrying five transgenes: Tg[*elavl3:H2B-GCaMP6s*]^jf^^5^ from Misha Ahrens^45^ in Casper background^62^, Tg[*Th1-*Gal4] AB line from Jiu-lin Du^43^, Tg[*UAS-NTR-mCherry*] line from Michael Parsons^63^ and Tg[*Th2-Gal4-VP16,UAS-E1b-NTR-mCherry*] AB line from Adam Douglass^32^.

Two-photon imaging experiments were conducted with zebrafish larvae obtained from five adult nacre cousin-pairs all carrying the five transgenes from four different sources. Since the larvae used for two-photon imaging were not purely homozygous in all the five transgenes, they were subjected to stringent fluorescence screening with 10-20% selection rates with highest-uniform expression of all transgenes. This necessitated the collection of thousands of fertilized embryos for each batch of experiments and ended with brain imaging of less than ten 7-8 dpf larvae. Collected embryos were screened under a Leica MZ FLIII stereo microscope on 4 dpf by anaesthetizing with Tricaine (∼0.16 mg/ml). Larvae expressing weak or no *Elavl3:H2B-GCaMP6s* (brain-wide green fluorescence) were sorted out into a separate dish. The GCaMP positive larvae were sorted into 5 groups based on red (mCherry) fluorescence visualized at 40-60x magnification: 1) not visible, 2) weakly/sparsely expressed, 3) bright Th1-mCherry, characterized by necklace pattern at the base of the dorsal hindbrain, 4) bright Th2-mCherry with linked expression in the eye-lens and 5) both Th1-& Th2-mCherry. Larvae with edemas and other developmental defects such as deflated swim bladder or bent spines were excluded. Finally, we obtained 20 to 50 larvae per week (total n=308) with robust-uniform expression of Th1-mCherry and Th2-mCherry. In case, too few larvae were obtained for behavioral experiments, those without *Tg[elavl3:H2B-GCaMP6s*] were subjected to the same screening but not used for brain imaging.

For behavioral experiments (shown in Fig. 2), a separate set of purely homozygous transgenic larvae (n=334) were obtained in subsequent generations from adult nacre sibling/cousin-pairs homozygous in all the four mCherry-associated transgenes: Tg[*Th1-Gal4: UAS-NTR-mCherry]* and Tg[*Th2-Gal4-VP16: UAS-E1b-NTR-mCherry*]. During fluorescence screening on 4 dpf, very few larvae (<10%) were discarded due to visibly weaker mCherry expression. However, a significant fraction of larvae had to be discarded due to morphological or developmental defects.

For behavioral experiments to investigate the loss of only Th1-expressing neurons, the parents were homozygous in Tg[*Th1-Gal4: UAS-NTR-mCherry*] and heterozygous in Tg[*elavl3:H2B-GCaMP6s*].

These experiments spanned three years and multiple (>4) zebrafish generations, addressing challenges of high mortality rates in nacre lines homozygous for multiple transgenes.

### Treatment with the pro-drug Metronidazole (MTZ)

On 4 dpf (evening) after fluorescence screening, zebrafish larvae expressing *Tg[elavl3:H2B-GCaMP6s*; *Th1-Gal4: UAS-NTR-mCherry; Th2-Gal4-VP16: UAS-E1b-NTR-mCherry]* were split into three groups by treating with: (A) 0.1% DMSO, (B) 5 mM MTZ with 0.1% DMSO and (C) 2.5 mM MTZ with 0.1% DMSO. A fourth group without any red fluorescence, was treated with 5 mM MTZ to serve as MTZ control. All larvae were kept in 10 cm Petri dishes in a 50 mL volume of E3 medium. All the solutions were replaced with fresh MTZ or DMSO solutions after ∼20 hours. On 6 dpf, ∼40 hours since the onset of drug/vehicle treatment, the MTZ/DMSO solution in all dishes were replaced by fish-system water for recovery. Pilot experiments carried with recovery in fresh E3 medium instead of fish-system water showed high mortality of MTZ-treated larvae, possibly due to significant wipeout of gut-microbiome^64^. Thus, 6 dpf onwards, E3 medium was only used for a few hours during two-photon imaging and acclimatization. Separate batches of similar experiments were also conducted using larvae without Tg[*Th2-Gal4-VP16: UAS-E1b-NTR-mCherry]* to investigate only the loss of Th1-expressing neurons.

### Free-swimming behavioral recording

After at least 24 hours of recovery in fish system water following 40 hours of MTZ/DMSO treatment, the larvae were transferred to a behavioral arena constructed of 6”x9” borosilicate glass bottom plate and laser-cut white acrylic walls, which were painted black on the top. It had 24 square wells each with 35mm width and 6 mm depth, filled with 7.5 mL of fish-system water. The arena was placed inside a Styrofoam box kept on top of a white LED panel (4500K). Two white A4 sheets of paper were placed between the plate and the box to function as a diffuser. An iPad Air 2020 (4^th^ generation) was placed on top of the lid (with a hole) for slow-motion HD (1920 x 1080) video recording at ∼240 frames per second. Some heat was generated by the white LED panel below the Styrofoam box which helped maintain a temperature of 25–26.5°C inside the recording box. The room temperature was 22-24°C and monitored using SensorPush through a smartphone app.

Just after placing zebrafish larvae inside individual wells, they were checked for startle response (gentle tap on the Styrofoam box). Larvae which did not respond to startle were removed from the behavioral arena and replaced by other larvae, followed by a second check for startle response. After ensuring all larvae were startle-responsive and that there were no debris in any of the wells, the lid was closed, and the iPad was placed on the lid. The nacre larvae (eyes) could be clearly seen on the large iPad screen and digitally zoomed to confirm that the focusing and exposure was suitable before starting the recording. The larvae were then acclimatized for ∼20 minutes prior to 15 minutes of video recording with locked camera exposure and focus. The iPad camera showed negligible barrel/pincushion distortion across the entire field of view. The 6 mm depth of wells ensured minimal parallax shadow.

Following the behavioral recording, larvae were handled and maintained individually in 6-well plates in the order of their position in the behavioral arena. Behavioral recording was performed on both 7 dpf and 8 dpf. No feeding (paramecium) was done until behavioral recording was complete, and larvae were transferred to the 6-well plates. No feeding was done until the night of 7 dpf even if the behavioral recording was completed earlier.

### Free-swimming behavioral tracking

The movie recorded using iPad in .MOV format was converted into .mp4 format (not required for Windows 11) with minimal loss of signal quality using ffmpeg with the following code executed through windows command line: *for %a in ("*.MOV") do ffmpeg -hwaccel cuda -i "%a" -c:v libx264 -an -crf 7 -filter:v "crop=in_w-20:in_h-220" -vframes 216240 "%∼na.mp4"*

The script crops out 220 rows from top and bottom, and 20 columns from left and right. It extracts the first 216240 frames, while the movie might contain more frames as the recording was manually stopped and thus had extra frames. The videos had a resolution of ∼0.14 mm/pixel, which typically represented the eyes of the nacre larvae as a cluster of 9-15 dark pixels against the background, which was estimated from the maximum projection of first and last 1-minute video frames. This background image was also used to detect the square arena for each larva. Using Otsu’s threshold with a multiplier of 1.2, from a gaussian smoothened image of the background we were able to identify all the black-painted boundaries to assign 24 arenas corresponding to each well.

For each of the 24 arenas, a custom segmentation algorithm isolated the pixels corresponding to eyes of the larva. The region close to the walls was susceptible to higher false positives/negatives, which was significantly reduced using appropriate logical and morphological filtering. Our algorithm worked flawlessly with pigmented larvae and juvenile larvae (not presented in this paper) which had significantly more dark pixels for tracking. The detected eye-cluster corresponding to each larva was saved as sparse matrices into a mat file along with several parameters such as centroid, area, edges, and signal-to-noise ratio. This program also generated a heatmap of time spent at each point in the arena, which allowed visual inspection before calculation of various parameters: (i) total distance travelled in 15 minutes, (ii) root-mean square speed, (iii) percentage of arena explored, (iv) number of bouts per second, (v) time interval between consecutive high-bouts, (vi) average distance moved in each bout, (vii) percentage duration of active swimming and (viii) percentage of time spent close to the walls vs. away from the walls (thigmotaxis). The boundary for thigmotaxis was 6.75 mm from the walls on all sides, which is 1.5 times the approximate larval body length (4.5 mm).

Statistical analysis was performed with ordinary one-way ANOVA followed by Tukey’s multiple comparison test using GraphPad Prism.

### Sample preparation for two-photon imaging

Zebrafish larvae were randomly selected for two-photon imaging on 7/8 dpf, from six-well plates where they were maintained systematically after behavioral recording. A small drop (∼100 µl) containing larva was placed at the center of a clean Borosil petri dish. Water surrounding the larva was carefully pipetted out while ensuring that the larva was not injured by the pipette tip (200 µl). Immediately, ∼70 µl of 2% low melting agarose (in 1x E3 buffer) maintained at ∼45°C was added on to the larva. The larva was immediately aligned upright using a soft plastic tip before the agarose solidified. A sharp knife was used to free the tail by cutting out solidified agarose below the swim bladder of the larva. This not only allowed the investigation of tailswing behavior, but also gauge the health of the larvae, few of which could be injured by the mounting procedure. It was ensured that mounted larvae showed tail-reflex when gently prodded by a thin plastic tip. Each larva was acclimatized for 1 to 2 hours in the tail-free mounted state in dark microscope room (maintained at 24°C) before brain imaging. Additionally, before the timelapse imaging for 24 minutes, all larvae were subjected to several minutes of two-photon imaging for acquisition of high-resolution Z-stacks and orientational adjustments. During this stage if the larva did not show any tail movements, the imaging would be discontinued. Similarly, at any point during the imaging, if brainwide GCaMP levels suddenly became saturated (which is typically indicative of death or injury), the imaging was discontinued (or such data were discarded later).

Pilot experiments indicated that most MTZ-treated larvae (but not controls) do not survive the mounting procedure on 8 dpf and there was not enough time on 7 dpf to image more than ∼5 larvae. Feeding larvae (both control and ablated groups) on 8 dpf was found to significantly improve the health and survivability of 8 dpf larvae for 2-photon imaging.

### Identification of CA-deficient larvae

We used a 35% cut-off for arena exploration to identify CA deficient larvae (not known during brain imaging). Only 3.5% (n=8) of all Th1&2 control larvae (n=228) explored less than 35% of the arena. To get a similar distribution of control larvae using other adversely affected parameters, for instance, total distance traveled, we would have to use a cut-off of 1.6 m, which drastically reduced the percentage of impaired CA-deficient larvae. Thus, we concluded that “arena exploration” was the most sensitive parameter affected by loss of CA neurons, which is not surprising considering dopamine’s role in motivation. With this parameter and 35% cut-off, we found nine CA-deficient larvae with timelapse two-photon data out of a total of thirty recorded Th1&2 larvae treated with 5 mM MTZ.

This significantly skewed distribution of CA-deficient larvae with brain imaging data does not reflect their free-swimming distribution as we only imaged larvae which were seemingly healthy and responsive after hours of acclimatization mounted in low-melting agarose until being subjected to functional brain imaging.

### Live brain imaging simultaneously with motion tracking

We custom-built a two-photon microscope using a Thorlabs galvo-resonant scan module and integrated it with dual Coherent femtosecond lasers (Axon 920 nm and Axon 1064 nm) and a Nikon 16× water-immersion objective (NA 0.8), all within a budget of $250,000. The system enabled simultaneous two-color imaging via spectral separation, with acousto-optic modulation of laser power across Z-depth to enhance signal-to-noise ratio in ventral brain regions. The focused laser spot was stretched along Z-axis by underfilling of the objective back aperture. The head of zebrafish larvae were consistently made to point towards the left and be parallel to the X-axis (horizontal), by manually rotating the stage. The stage also allowed adjusting the tip-tilt (<5°) of the sample, which was useful to maintain a consistent orientation for the brain. The complete list of custom components and system configuration can be made available upon reasonable request.

A monochrome CMOS camera with ½” sensor (FLIR Chameleon) was used to record the tail behavior from the bottom of the sample stage. To achieve an overall 1× magnification at the camera sensor, we demagnified the image from the microscope objective (Olympus Plan N 4×, NA 0.10) using a custom tube lens configuration. Pixel binning and cropping were used to boost the frame rates to ∼200 fps, while the movie was recorded using “mjpeg” compression in avi format. The illumination for tail recording comprised of three infrared lamps at ∼800 nm placed strategically around the sample stage.

For functional imaging, we acquired whole-brain volumetric data in the green channel at 1 Hz for 1401 frames (∼24 minutes). The imaging volume covered a 1000 × 500 µm field-of-view (1024 × 512 pixels) with a depth of 270 µm sampled by 28 Z-planes (10 µm step-size) and 3 flyback frames. A polygon mask was drawn (at the very beginning) on the live image to protect the eyes of zebrafish larvae from 920 nm laser, along with a custom non-linear laser-power profile. Following the live imaging, a Z-stack was acquired at the same XY resolution (0.977 µm/pixel) but increased Z-resolution (2 µm per step) using conventional staircase mode instead of high-speed continuous ramping which was used for functional imaging.

### Extraction of spatiotemporal component of neurons from two-photon data

Two-photon timelapse data was saved by ThorImage®LS as a 3D stack, which was converted to a 4D hyperlapse using a macro in ImageJ with appropriate parameters, discarding the 1^st^ volume. It was then saved as a single tiff file using the bioformats plugin in ImageJ using the same macro, and analyzed using suite2p (version 0.7.5) for preliminary extraction of the spatiotemporal components of the neurons. We analyzed 57 larvae out of approximately 70 recordings. Larvae were excluded from analysis due to unsuccessful motion correction, Z-drift, absence of tailswing activity during two-photon imaging, or GCaMP saturation at any point during the timelapse (from abrupt increase in Ca++ due to photothermal damage). Of the analyzed larvae, 43 expressed all five transgenes. Within this group, 24 larvae were treated with 5 mM metronidazole (MTZ), 6 with 2.5 mM MTZ, and 13 served as DMSO controls. The remaining larvae consisted of two control groups: 7 larvae lacking red fluorescence (treated with 5 mM MTZ as sibling controls) and 7 larvae expressing TH1-Gal4 but not TH2-Gal4.

A significant amount of false positive neurons and activity was detected by suite2p despite several attempts at optimization of various parameters and training of the in-built CNN classifier. CaImAn also had a similar issue, but suite2p was preferred mainly due to its user-friendliness and computational speed. To remove false positive neurons and activity, a custom MATLAB program was used which could not only remove false positive neruons based on morphology and activity, but could also merge neurons which had been wrongly split. The activity of all neurons in the larvae were normalized between 0 and 1, following which dynamic baseline subtraction (using moving window of 2 minutes) was applied to the activity of each neuron. For few randomly selected neurons, the extracted temporal component was verified by visual inspection of the movie. This neuronal activity was summed across time for each neuron, followed by z-score normalization for comparison of cumulative ΔF/F across larvae.

To estimate neuronal firing rate corresponding to the fluctuations in GCaMP fluorescence of each neuron, neuronal spikes were inferred from suite2p’s raw fluorescence traces using CNMF’s^65^ constrained FOOPSI AR1 model (solved by CVX ver.2.1). The total number of peaks in the deconvoluted spike train was divided by the total imaging duration to calculate the firing rate of each neuron. The spatiotemporal component of neurons were saved in .mat files for further processing and analysis.

### Creation of custom reference template and masks for anatomical registration

Since different brains were recorded at varying orientations, it was crucial to align them with a reference brain for comparison of neural activity. We aligned *Elavl3-H2B* reference templates from both Z-Brain^41^ and mapZebrain^42^ atlas to one of our own recorded *Elavl3-H2B* z-stack by using ANTsPy implementing *SynAggro* transformation. All our other recorded brains were also aligned to this reference brain. We found this non-conventional method of anatomical registration to be more accurate than aligning all recorded brains directly to Z-brain, which was volumetrically slightly smaller than our recorded 7/8 dpf brains (Extended Data Fig. 1). This was not surprising because z-brain is based on anti-ERK staining average of 6 dpf nacre larvae, which in turn is based on the classical neuroanatomical studies of 3 dpf and 5 dpf zebrafish brain^66^.

The transformation matrices obtained from the alignment of *Elavl3-H2B* templates were used to transform 294 anatomical masks from Z-brain, 114 anatomical masks from mapZebrain and 296 HCR/transgenic markers from mapZebrain. Additionally, we sub-divided each of these masks based on glutamatergic, GABAergic, and cholinergic signaling, generating a total of 2158 masks. We built a custom mask for Mesencephalic Locomotor Region (MLR) and another mask from the loss of both Th1 and Th2 expressing neurons.

### Quantification of CA-neuron-loss due to chemogenetic ablation

In addition to the timelapse acquisition, a z-stack was acquired to map the distribution of CA neurons across the entire brain with a resolution of 0.65 × 0.65 × 2 µm in green and red channels. The imaging depth was increased to 400 µm from the 270 µm used for functional imaging. Prior to alignment using ANTsPy, a z-offset of 10 µm was applied to the red z-stack to account for chromatic aberration of 1064 nm laser (red channel) with respect to 920 nm laser (green channel). This z-offset was estimated independently by imaging neurons co-expressing GFP and mCherry (not presented here). The actual value of chromatic aberration varied with imaging depth, and 10 µm was considered a good approximation. The acquired green and red z-stacks were aligned to the reference brain followed by calculation of the overlap between the aligned red channel and twelve anatomical mask-composites from z-brain atlas (**Supplementary Table 1**).

For estimation of the overlap, initial threshold of 0.2 was applied for counting voxels, following 0-1 normalization of each z-stack. The threshold value was increased or decreased iteratively in subsequent steps by identifying the outliers in each round. Otsu’s threshold was not used because it was found to be very sensitive to the noisy background of MTZ-treated larvae. The percentage loss of CA neurons was calculated as 100*(a-b)/a, where a and b are the TH-voxel counts in control and MTZ-treated larvae, respectively. These results were then plotted using GraphPad prism. Statistical significance was determined using Wilcoxon rank-sum test and Permutation test, whichever yielded higher p-value.

### Assigning neurons to anatomical groups

The weighted centre-of-mass of neurons extracted from our MATLAB version of suite2p-CNMF, was aligned to the reference brain using ANTsPy’s ‘apply_to_point’ function based on the transformation matrix generated from the alignment of z-stack (2 µm z-steps) corresponding to the timelapse-volume (10 µm z-steps). A z-offset of 5 µm (half of the z-step-size) was added to each neuron’s z-position prior to alignment to account for the steady motion of the microscope objective during the timelapse acquisition. This was an exclusive feature of our microscope that it did not halt intermittently at every z-step which is typical for “staircase mode” on most microscopes, and thus enabled similar frame rates as with remote focussing.

The aligned coordinates were used to construct a cube of 75 voxels (5 × 5 × 6 µm) representing each neuron, which was then mapped into a stack of 500 × 1000 × 200 voxels, corresponding to a size of 500 µm × 1000 µm × 400 µm. The resulting stack of each neuron was then checked for overlap with the binary masks we generated from z-brain and mapZebrain, thus creating a 2D overlap matrix for each larval brain. Neurons with at least 10 voxels overlapping with an anatomical mask were assigned to that anatomical group.

The z-scored cumulative ΔF/F for each neuron belonging to certain anatomical group was summed up for comparison with other larvae, while the number of spikes was averaged to compare firing rates. Anatomical regions were defined as part of hindbrain, midbrain and forebrain if more than half of their volume was contained within one of the three large brain regions.

### Voxel-based neural activity comparison

In addition to the conventional approach of analyzing neuronal activity based on anatomical groups, we decided to pursue a parallel approach which would not be bound by known anatomical information and could possibly identify novel brain regions. For this, the aligned coordinates of each neuron were downsampled from 1 × 1 × 2 µm to 12 × 12 × 12 µm. This enabled the spatial mapping of cumulative ΔF/F, number of active neurons and average firing rate as 12 µm voxels, generating a downsampled spatial heatmap of size 42 × 84 × 32 voxels.

The value of cumulative ΔF/F, number of active neurons and average firing rate corresponding to each heatmap-voxel was statistically compared between control and impaired larval groups using Wilcoxon rank-sum test (p<0.05). This allowed us to identify which voxels were significantly altered by the loss of CA neurons. Interestingly, many of those statistically significant voxels could be mapped to known anatomical regions. Additionally, it highlighted the heterogeneity inside individual anatomical masks, particularly the telencephalon, which otherwise averaged out to be unaltered by CA loss.

### Extraction of tailswing behavioral data during 2p-imaging

Tail-swing behavior recorded during functional brain imaging was analyzed using a custom MATLAB program. This program uses GPU acceleration to skeletonize the tail, which works even if the tail appears blurry due to movements faster than the imaging frame rate (tested 100 fps to 200 fps). This program saves XY coordinates as sparse matrices, representing the skeletonized tail for each frame of the movie, between the user-specified start to stop frame. The tail is split into 8 segments based on the number of points (not length) representing the tail. This is followed by fitting a 3^rd^ order polynomial to each segment and calculating the angle it makes with the X-axis by fitting a line to the polynomial. For the analysis reported here, a single angle was derived by averaging the points from segments 4 to 7, followed by measuring the slope of the line connecting it to the user-selected basepoint. Any nan value for the angle was replaced by their nearest non-nan value, which was usually encountered for not being able to detect the tail in a certain frame due to poor signal-to-noise ratio for detection of the tail over the background. This was followed by dynamic baseline removal estimated for every 1-minute bidirectional window. This was necessary because the resting state of the tail could slightly change with time over long durations. A plot of maximum projection of tail movie every 15 seconds overlaid with detected tail-points allowed visual verification of the accuracy of tail tracking (**Supplementary Fig. 6**).

### Analysis of tailswing behavior

The tail-angle data was analyzed using a custom MATLAB program to identify all tail-swing events. The noise floor was calculated as the root-mean-square of the absolute value of all tail-angles. To be qualified as a tail-swing event it should have a peak value three times higher than the noise floor and minimum peak prominence higher than the noise floor. The onset and offset for each event were determined such that they did not overlap with adjacent events. If the gap between the offset of one event and the onset of next event was less than 2 seconds, then the events were merged.

Following the detection of tail events, struggle events were identified as those with bidirectional peak value ten times higher than the noise floor, which was approximately 45° for most larvae. The remaining events were classified as either left flip or right flip based on the sum of angles higher than three times the noise floor. For the right flip, the sum of positive angles was 20 times higher than the absolute sum of negative angles; while for the left flip, the absolute sum of negative angles was 20 times higher than the sum of positive angles. All remaining tail events were classified as “swim” without delving deeper into finer complexities.

### Correlating neuronal activity to tailswing behavior

The “start” and “stop” frames were manually identified from the tail tracking movies, which clearly showed the infrared laser scanning over the head of the larval zebrafish. The tailswing angle between the “start” and “stop” frames was then correlated with timelapse ΔF/F data from neurons by downsampling it to 1 Hz (same as neuronal timelapse data). We found that downsampling using MATLAB’s ***interp1*** function led to significant reduction of tail amplitude. Instead, we assigned the maximum angle, after removing outliers above 90^th^ percentile, as the tail angle corresponding to each neuronal timepoint.

The tail angle data was normalized and appended as an additional last row to the neuronal data matrix, which was previously normalized separately between 0 and 1 after z-score normalization across time. Hierarchical clustering was applied to the combined data matrix using pairwise distances computed via Spearman’s rank correlation (MATLAB’s pdist function). Spearman’s correlation was chosen for its non-parametric nature, which is better suited to neuronal data that often exhibit monotonic but non-linear relationships. It also provides robustness to outliers, which are common in neuronal fluorescence traces.

Clustering was performed using MATLAB’s linkage function with the average linkage criterion, generating a dendrogram representing the hierarchical structure. The clustering began with a predefined number of clusters (t = 3). Using MATLAB’s cluster function, each data point was assigned a cluster label. To identify the motor-associated cluster, the algorithm iteratively searched for the cluster containing the tail movement data (represented by the last row in the matrix). Stability of this cluster was ensured by verifying that its neuron count did not drop by more than 20% in any iteration. This process continued until a stable clustering configuration was reached. Neurons grouped with the tail movement trace were identified as part of the motor-associated cluster, while the remaining neurons were classified as non-motor. These neuronal groups were then mapped across 2158 brain regions for estimation of cumulative ΔF/F and average tail-neuron correlation.

### Connectivity Analysis

Spearman’s rank correlation was measured between neurons of one group (MLR, Tel crhb, etc.) with neurons in remaining groups. A threshold of 0.8 was then applied to detect neuron-pairs with remarkably high temporal correlation. The percentage of these neuron-pairs was used to define the connectivity between the corresponding region-pairs. The connected regions we identified are also true for lower threshold values but were not chosen to be as stringent as possible.

## Acknowledgements

We would like to sincerely thank Professor Allan Basbaum at UCSF for his invaluable feedback and insightful comments on this manuscript. His thoughtful suggestions greatly enhanced the quality of this work. We also thank Misha Ahrens, Adam Douglass, Michael Parsons, and Jiu-lin Du, as well as the Zebrafish International Resource Center (ZIRC), for generously providing transgenic zebrafish lines. We are grateful to Michael Munchua, Devin Wright, and Evan Lee for their assistance with fish care and maintenance. We also thank Mahendra Wagle for valuable research discussions and ongoing support, and Kushagra Bhardwaj for assistance with custom illustrations.

This work was supported by the U.S. Department of Defense (DOD) under award number W81XWH-18-1-0176, and by NIH grant R01NS120219 awarded to S.G. Microscope construction was supported by NIH R01 GM132500 (Equipment Supplement NOT-GM-20-013) awarded to S.G.

