## Extended Data Fig. 1 to 8 for "Multimodal sensory overload in dopamine-deficient larval zebrafish leads to paradoxical kinesia"

**Extended Data Figures**

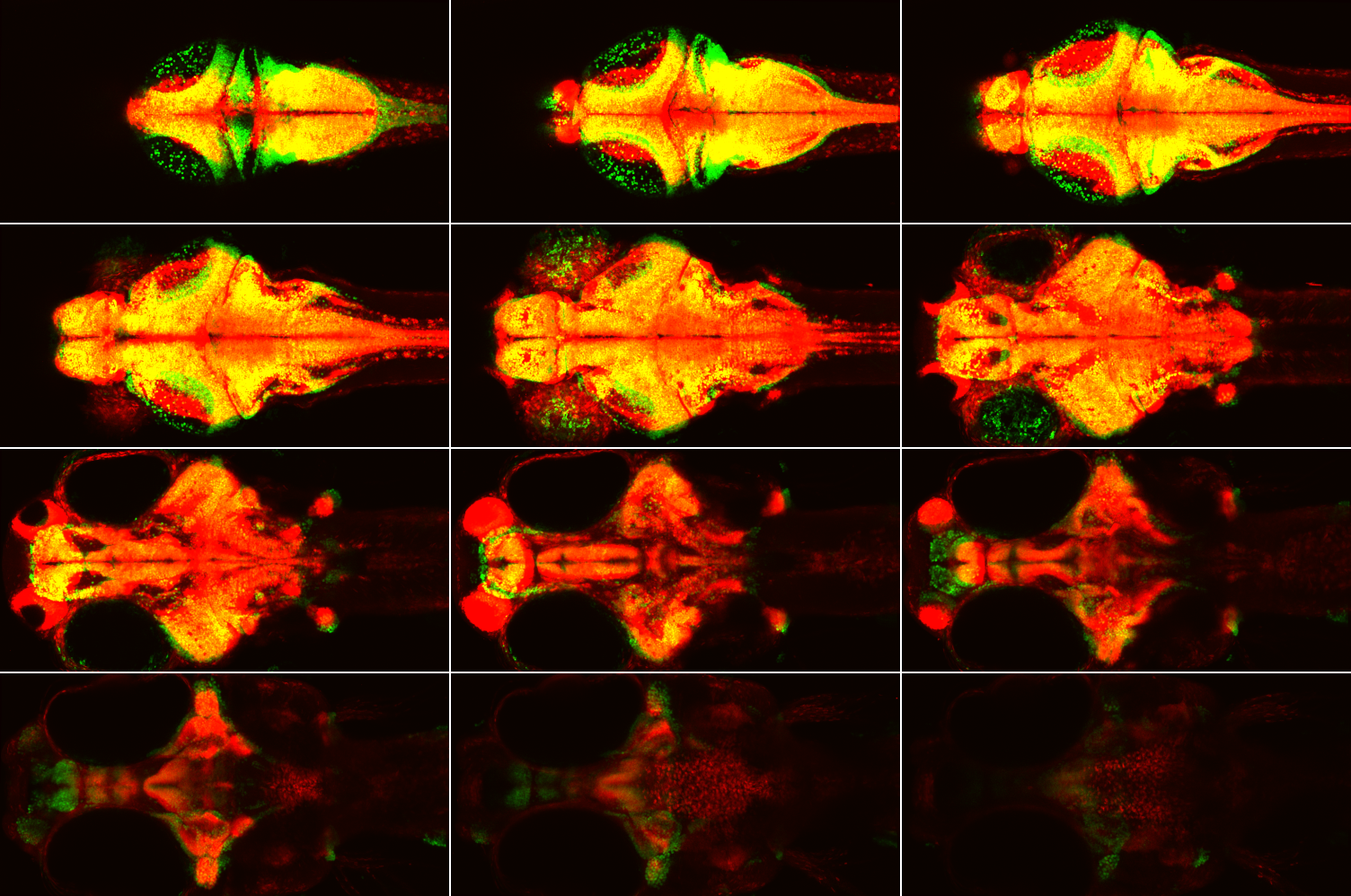

**Extended Data Fig. 1 | Smaller size of z-brain template compared to recorded brains.** Elavl3-H2B expression in 6 dpf Z-Brain template (red) and our 7 dpf reference template (green) aligned by eye-estimate to highlight the difference in brain size. The templates were resampled to have the same 1×1×2 µm resolution and resized/cropped to 1000×500×156 voxels. Each panel in the 4×3 montage is a 12-µm maximum z-projection. Field of view: 1 mm × 0.5 mm.

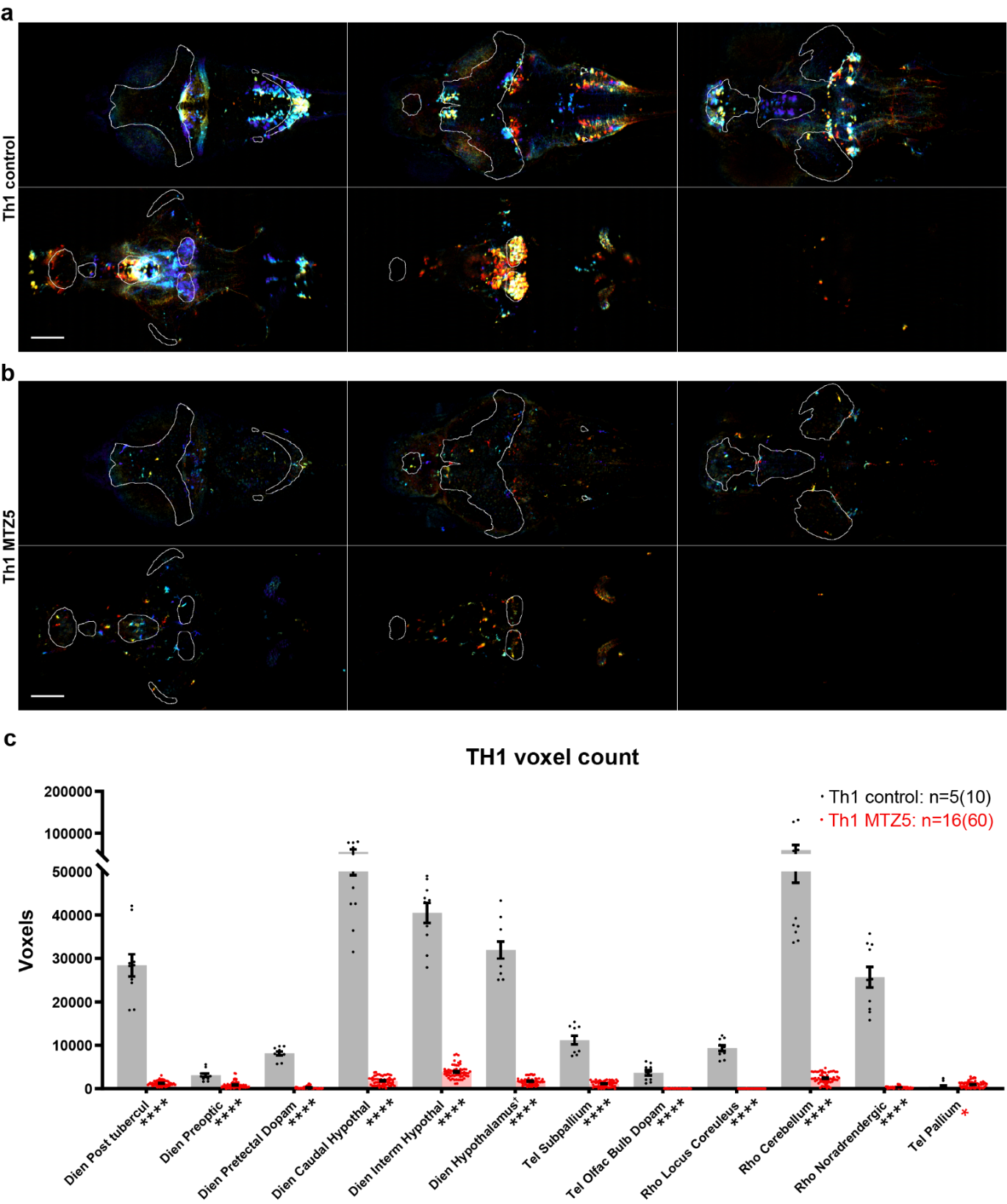

**Extended Data Fig. 2 | Chemogenetic ablation of TH1-Gal4 expressing neurons. (a)** Montage of depth-colored maximum intensity projections across 52 µm z-depths of two-photon imaging data from control **(b)** and MTZ-treated **(c)** larvae following ANTs registration. Images are divided into dorsal and ventral halves and depth-coded for better visualization. Both Z-stacks were processed with the same intensity ranges before depth coding. Anatomical regions are delineated by contours: telencephalon (white), diencephalon (grey), rhombencephalon (pink), and Z-Brain-defined dopaminergic regions (red). Scale bar: 100 µm; field of view: 1 mm × 0.5 mm. Quantification of Th1-Gal4 expressing neurons across 12 anatomical regions defined by the Z-Brain atlas. Bar graph shows mean voxel counts with individual data points overlaid (5 control larvae,16 MTZ-treated larvae). Asterisks adjacent to region-names indicate statistical significance (Wilcoxon rank-sum test and Permutation test: **** for p≤0.0001, * for p<0.05). Red asterisk indicates that control has lower value. †Caudal and Intermediate hypothalamus were removed from Hypothalamus.

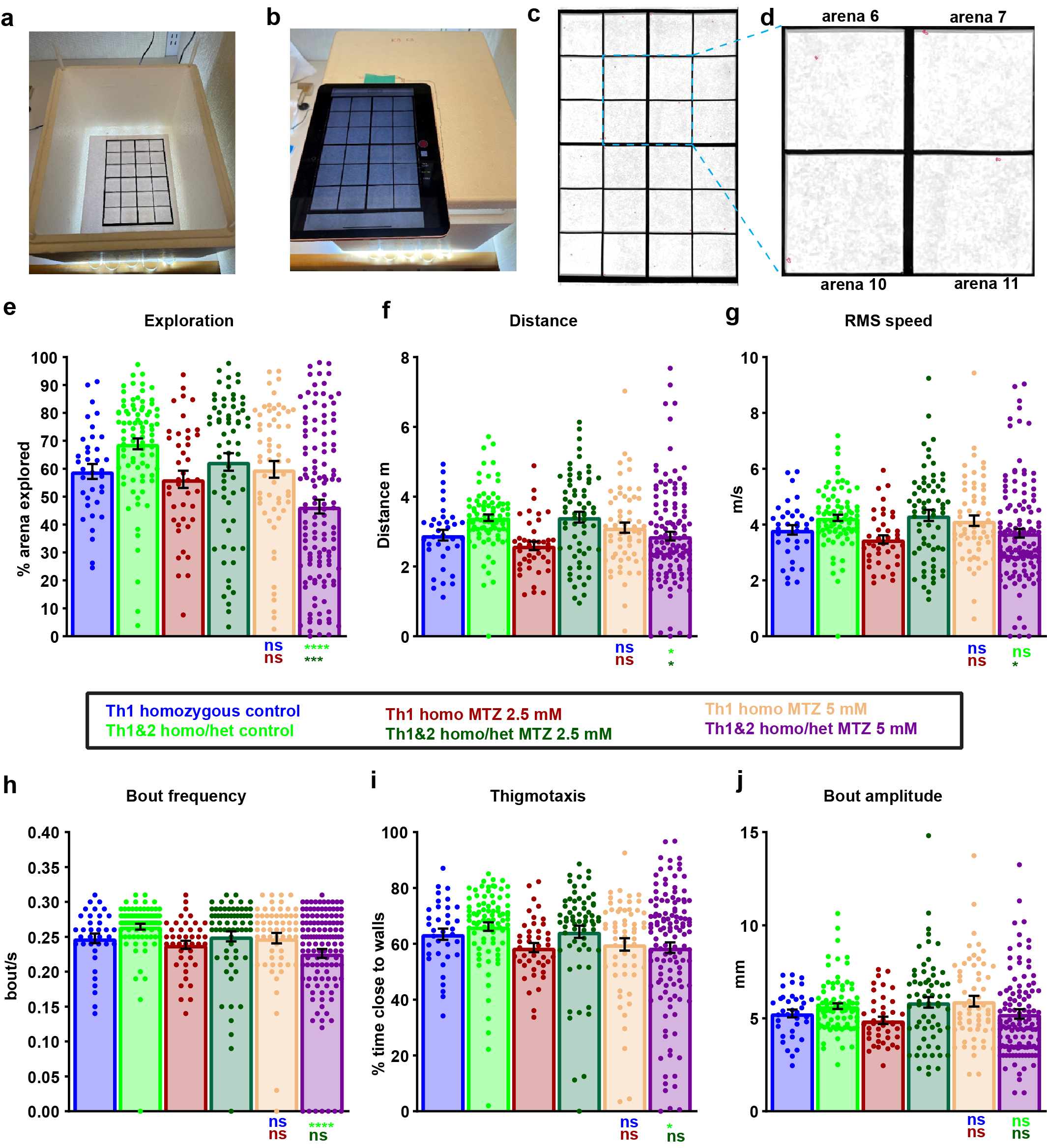

**Extended Data Fig. 3 | Behavioral analysis of free-swimming Th1 and Th1&2 larvae. (a,b)** Custom behavioral setup comprising a Styrofoam box, 6"×9" Borosil glass plate, laser-cut acrylic wells, 1 ml pipette tips (for height adjustment), and an iPad4 2020. Individual arenas (35 x 35 mm) contain 7.5 mL water each, with 24 arenas total. **(c,d)** Representative tracking output from Matlab-based program. **(c)** Single movie frame showing detected larvae outlined in red across all arenas. **(d)** Magnified view of four adjacent arenas highlighting individual larval detection. **(e-j)** Quantification of behavioral parameters across experimental groups: Th1 DMSO control (n=36), Th1&2 DMSO control (n=84), Th1 2.5 mM MTZ (n=42), Th1&2 2.5 mM MTZ (n=65), Th1 5 mM MTZ (n=60), and Th1&2 5 mM MTZ (n=126). “Th1&2” larvae represent the brightest ~30% of homozygous and heterozygous carriers expressing all three transgenes (Th1-Gal4, Th2-Gal4-VP16, and UAS-NTR-mCherry). Parameters analyzed: **(e)** percentage of arena explored, **(f)** total distance traveled, **(g)** root-mean-square speed, **(h)** bout frequency, **(i)** thigmotaxis (time spent near walls), and **(j)** bout amplitude (distance per bout). 5 mM MTZ treatment significantly reduced most behavioral parameters compared to DMSO-treated siblings and non-NTR-expressing MTZ controls, except bout amplitude which remained unchanged. Statistical significance determined by ordinary one-way ANOVA with Tukey’s multiple comparison test (**** p<0.0001, *** p<0.001, * p<0.05, ns p≥0.05).

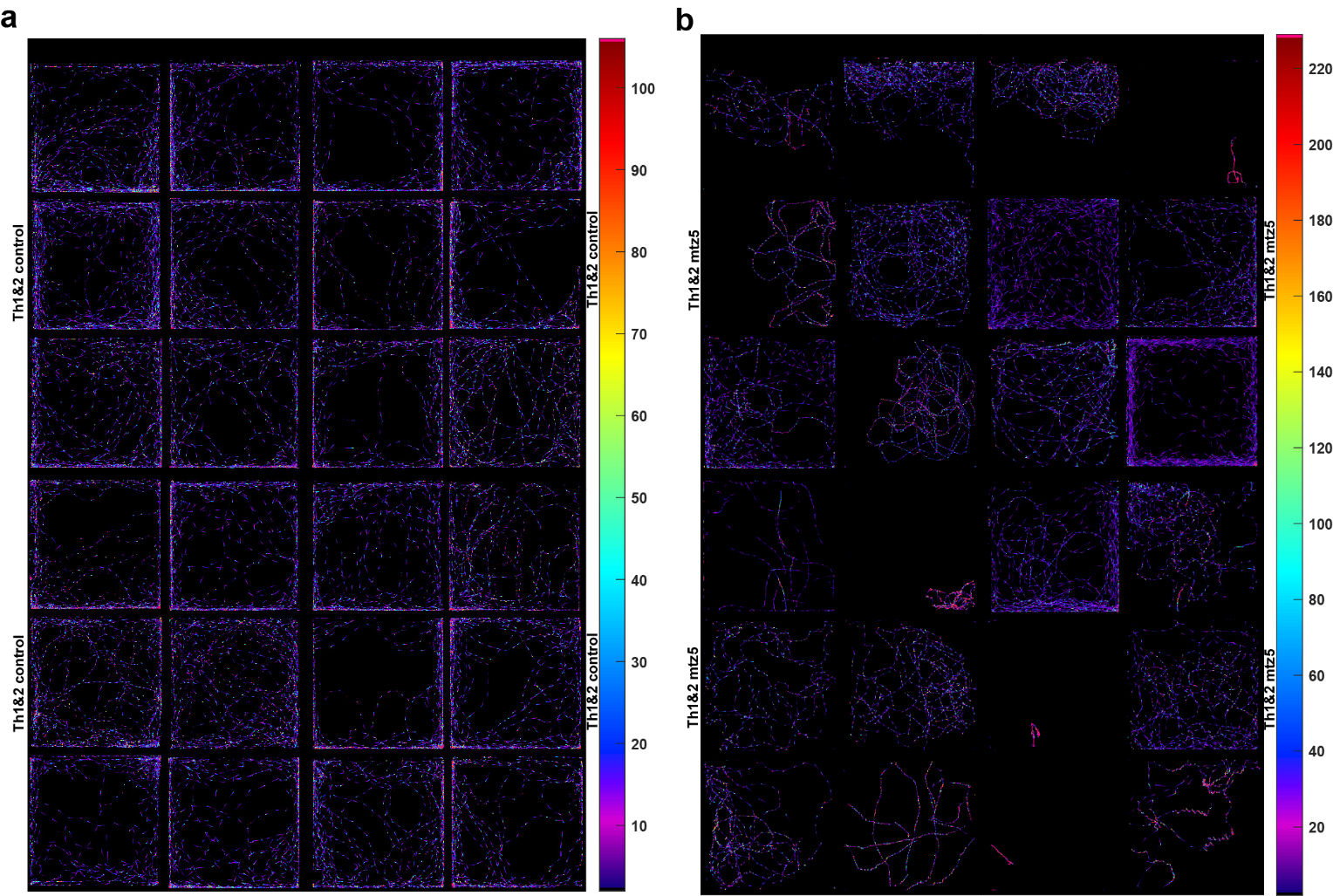

**Extended Data Fig. 4 | Spatial distribution analysis of control and CA-ablated larvae during free swimming. (a,b)** Heatmaps showing positional dwelling time during 15-minute recordings of **(a)** DMSO control larvae (n=24) and **(b)** Th1&2 ablated larvae (n=24). All larvae were homozygous for TH1-Gal4, TH2-Gal4, and UAS-NTR-mCherry transgenes. Color intensity indicates dwelling time at each pixel position, with red/pink representing longer durations. Note the expanded colorbar range (approximately doubled) for ablated larvae, reflecting increased positional dwelling time. Colorbar spans 2nd to 98th percentile of all non-zero pixel values.

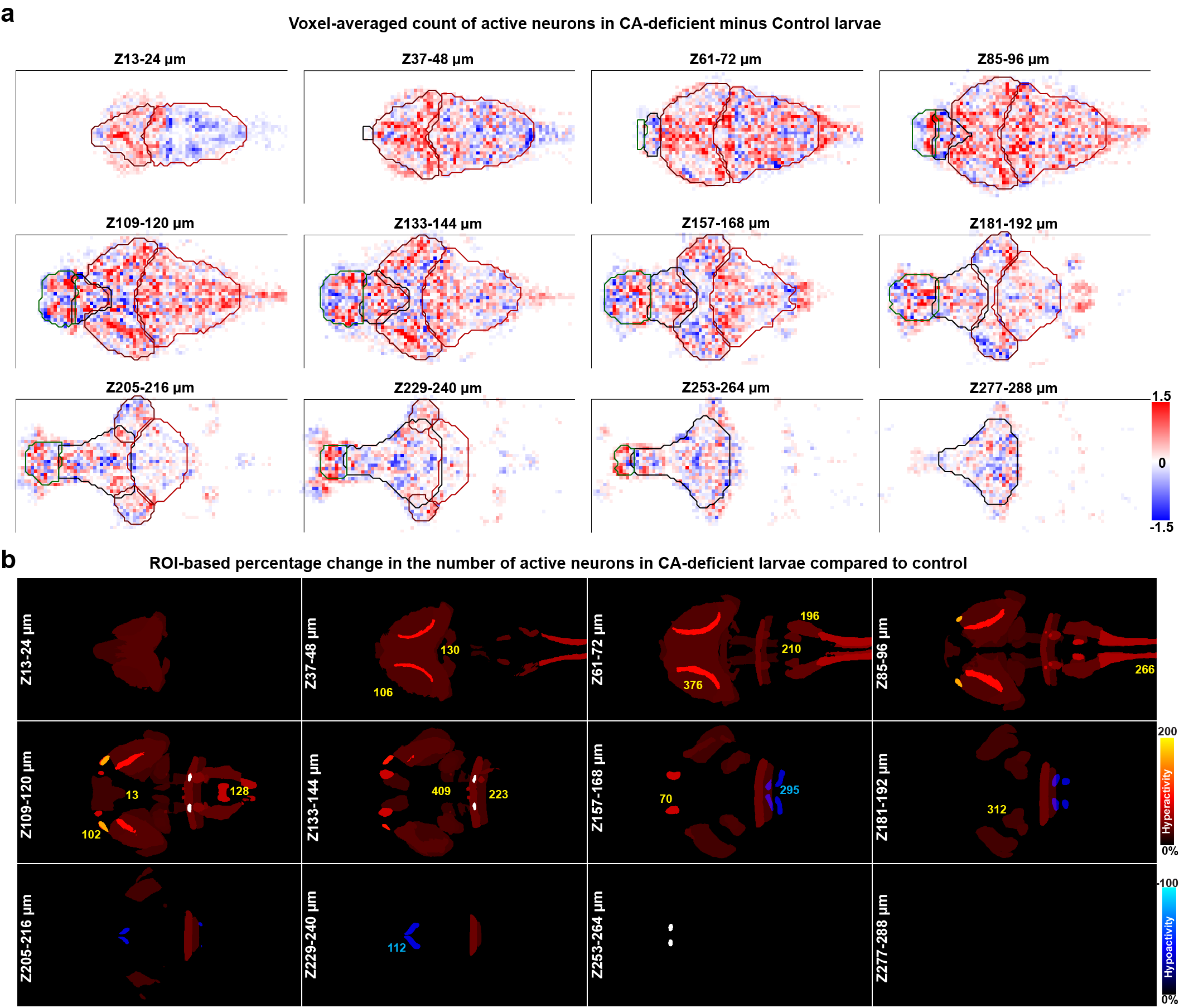

**Extended Data Fig. 5 | Altered number of active neurons in CA-Deficient Larval Zebrafish. (a)** Voxel-based comparison of active neuron-count revealed significant local variations between control and CA-deficient groups. Increased neuron count is indicated by red pixels, while reduced neuron count is indicated by blue pixels. The colorbar was saturated at ±3 to enhance contrast. **(b)** ROI-based view of anatomical regions showing altered neuronal activity. The regions are color-coded based on the percentage difference in firing rate between CA-deficient and control groups. Warm colors (black-red-yellow) indicate increased number of active neurons (>10%), while cool colors (black-blue-cyan) indicate reduced count of active neurons (<-10%) in the CA-deficient group compared to controls. Regions with activity ≥200% are shown in white. Each panel in the 4×3 montage is a 12-µm maximum z-projection, skipping every alternate 12-µm projection to reduce redundancy. **(a,b)** Field-of-view: 0.5 mm (vertical) × 1.0 mm (horizontal).

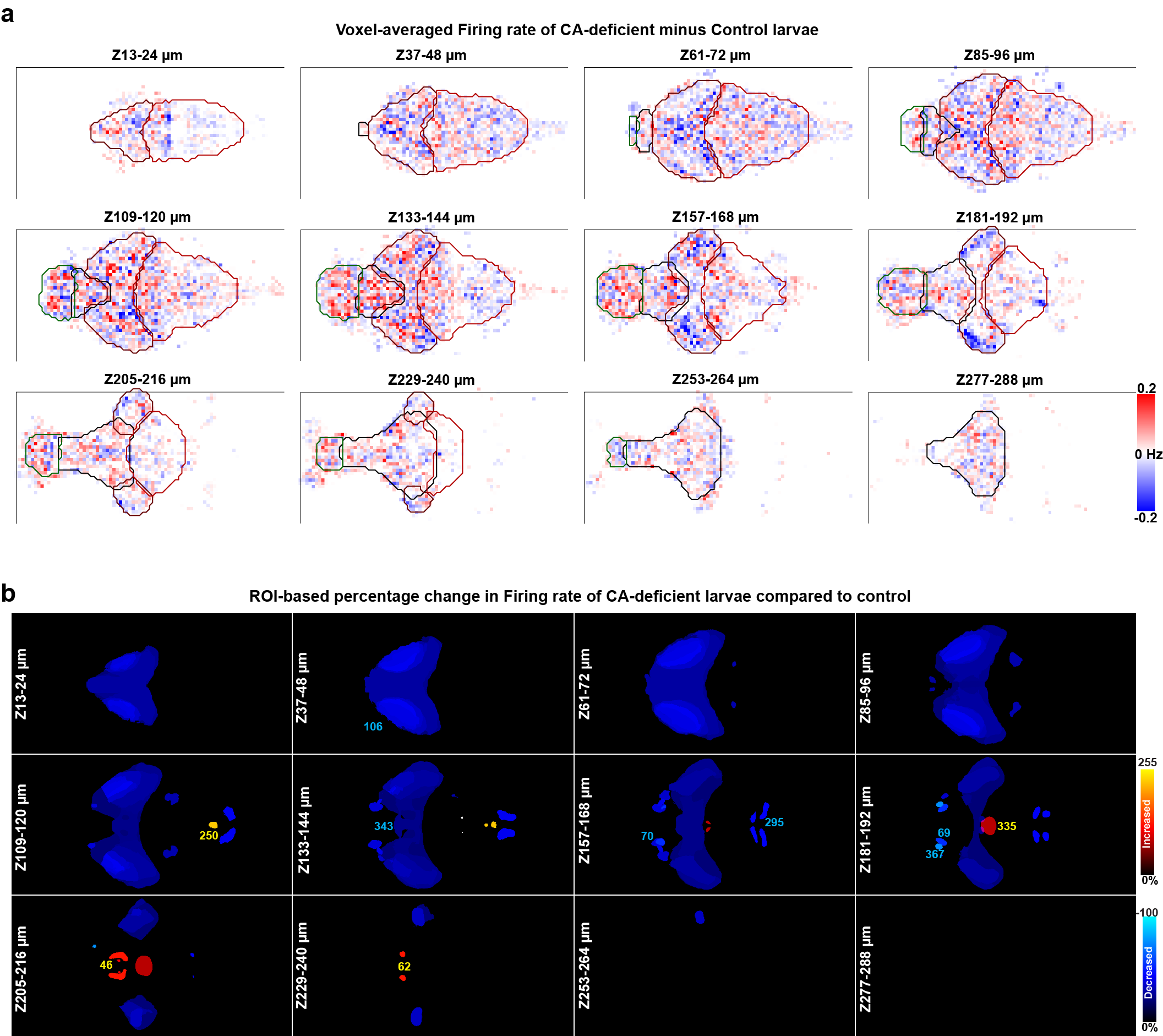

**Extended Data Fig. 6 | Altered Firing Rate in CA-Deficient Larval Zebrafish. (a)** Voxel-based comparison of firing rates revealed significant local variations between control and CA-deficient groups. Increased firing rate is indicated by red pixels, while reduced firing rate is indicated by blue pixels. The colorbar was saturated at ±0.2 Hz to enhance contrast. **(b)** ROI-based view of anatomical regions showing altered firing rate. The regions are color-coded based on the percentage difference in neural activity between CA-deficient and control groups. Warm colors (black-red-yellow) indicate increased firing rate (>10%), while cool colors (black-blue-cyan) indicate decreased firing rate (<-10%) in the CA-deficient group compared to controls. Regions with activity ≥200% are shown in white. Each panel in the 4 × 3 montage is a 12-µm maximum projection, skipping every alternate 12-µm projection to reduce redundancy. **(a,b)** Field-of-view: 0.5 mm (vertical) × 1.0 mm (horizontal).

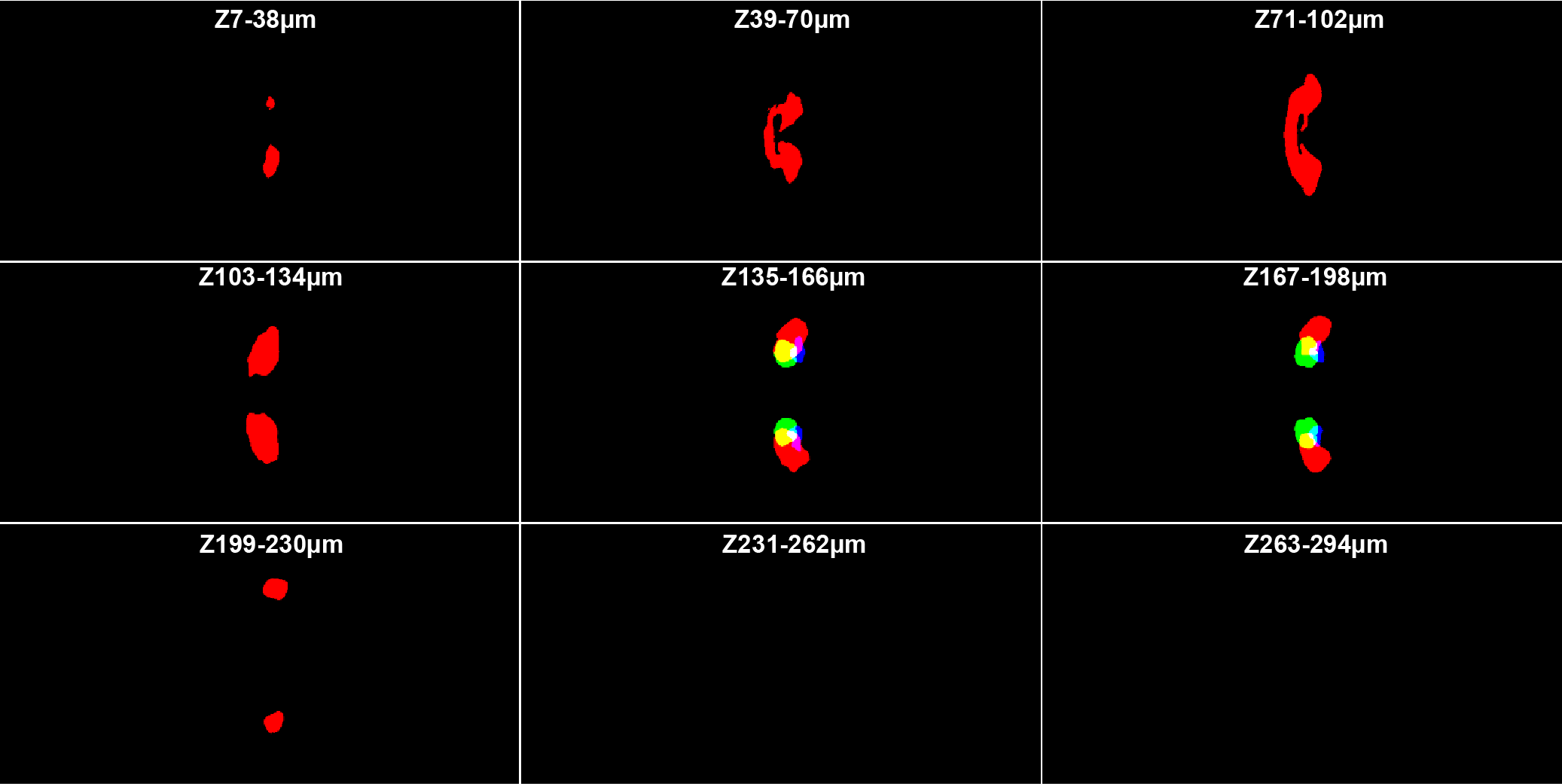

**Extended Data Fig. 7 | Proposed location of MLR (Mesencephalic Locomotor Region).** MLR is shown as a green mask overlaid with Rho Cerebelluar-Vglut2 enriched areas (#130, red mask) and Rho Locus Coeruleus (#183, blue mask). Each panel in the 3 x 3 montage is a 32 µm maximum z-projection.

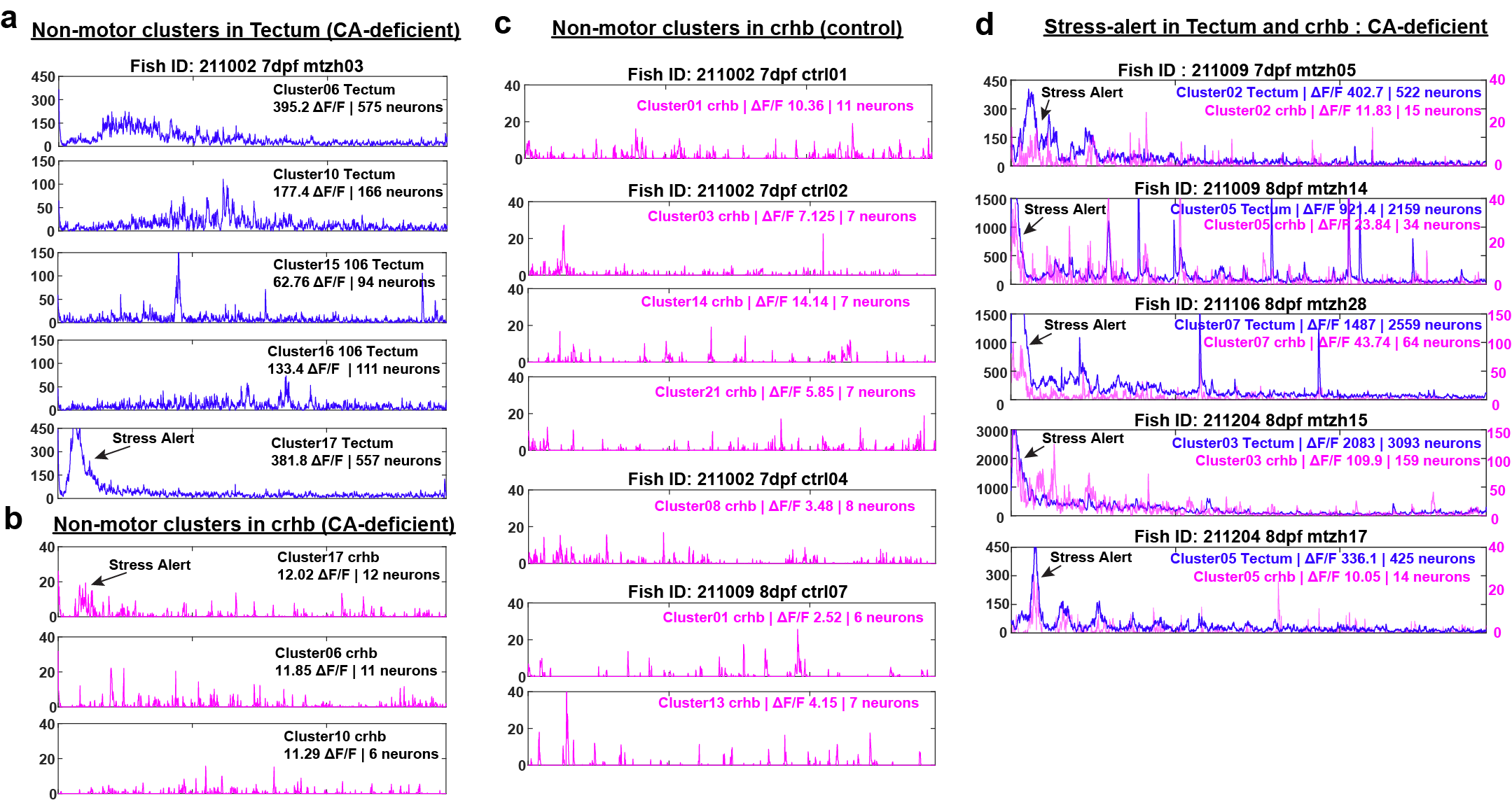

**Extended Data Fig. 8 | ΔF/F of neurons belonging to non-tail clusters in tectum and crhb.** Panel **(a)** shows the largest 5 of 20 non-motor clusters identified in the tectum of a representative CA-deficient larva (also shown in Fig. 6). Panel **(b)** shows all non-motor clusters identified in crhb-neurons of the same larva. Towards the beginning of 2p-imaging, a hump was observed in the ΔF/F of neurons belonging to cluster 17 of both tectum and crhb, as indicated by an arrow. **(d)** Similar humps were observed in five more CA-deficient larvae (out of 9) as indicated by an arrow in the summed ΔF/F of clusters 2, 5, 7, 3 and 5. **(c)** No such humps were observed in any non-motor crhb clusters across 10 of the 11 control larvae, of which four are shown. The number of non-motor neurons and their cumulative ΔF/F is shown in respective figure legends for tectum and crhb.
