## Supplementary Data Fig. 1 to 6 for "Multimodal sensory overload in dopamine-deficient larval zebrafish leads to paradoxical kinesia"

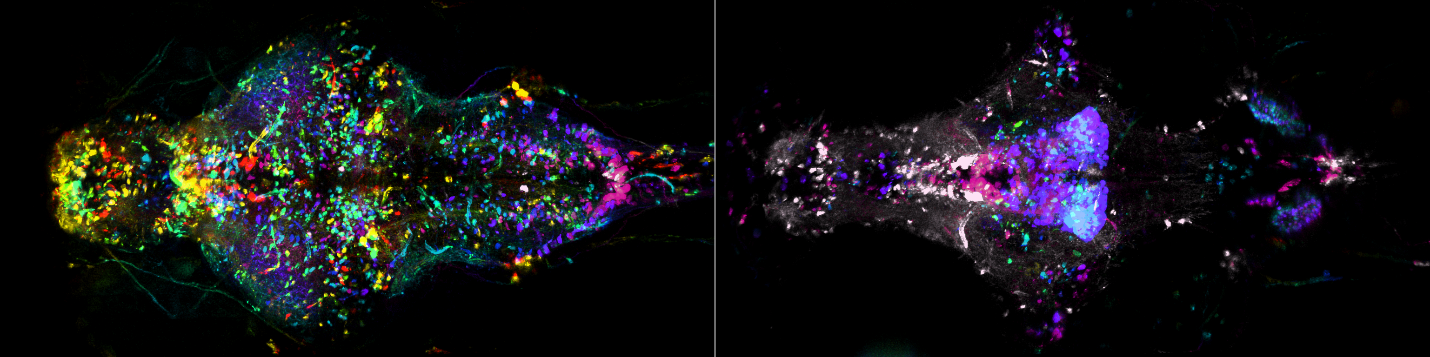

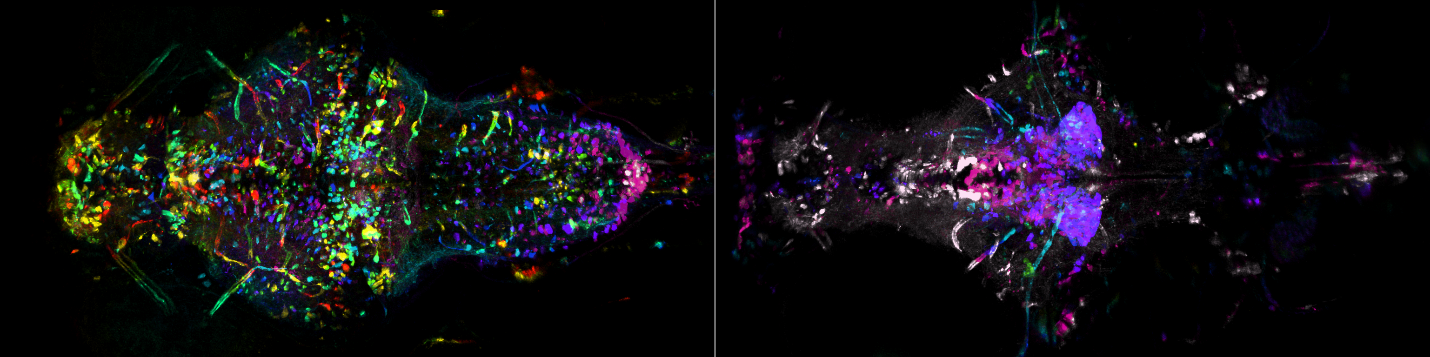

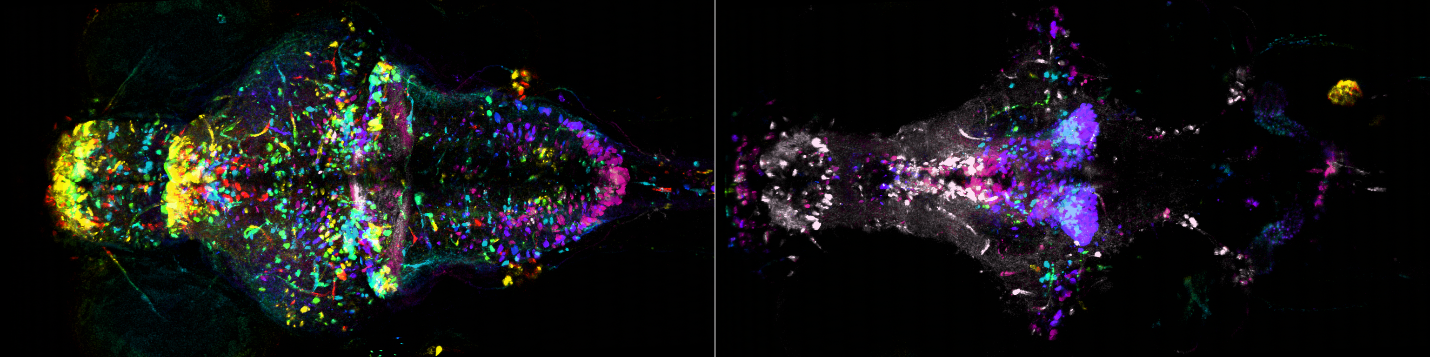

**Supplementary Fig. 1 | Spatial distribution of TH1 and TH2 neurons in control larvae.** Depth-colored maximum intensity projections showing UAS-NTR-mCherry expression driven by*Th1-Gal4* and *Th2-Gal4* in dorsal (left) and ventral (right) brain of 7-8 dpf larval zebrafish treated with 0.1% DMSO. Maximum projections were obtained across 200 µm for each half following registration using ANTs. Field-of-view: 0.5 mm (vertical) x 1.0 mm (horizontal).

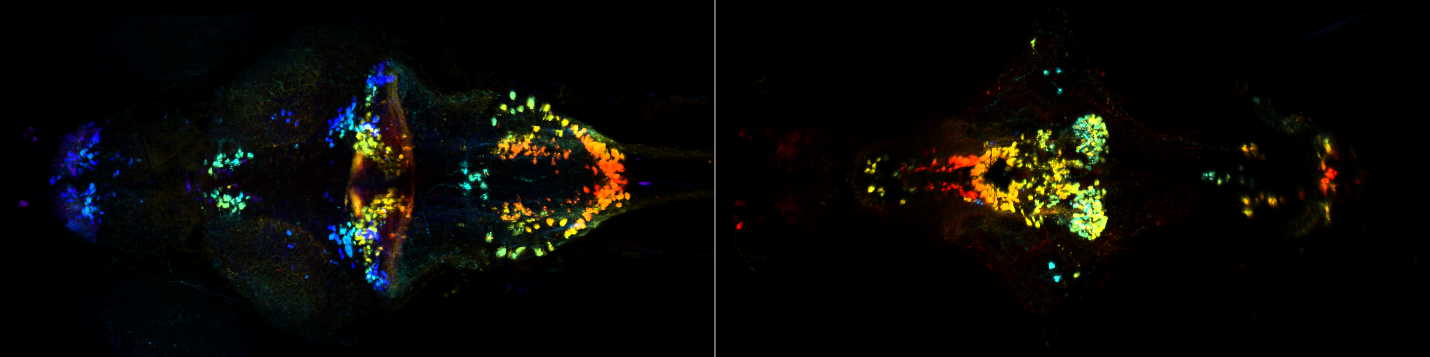

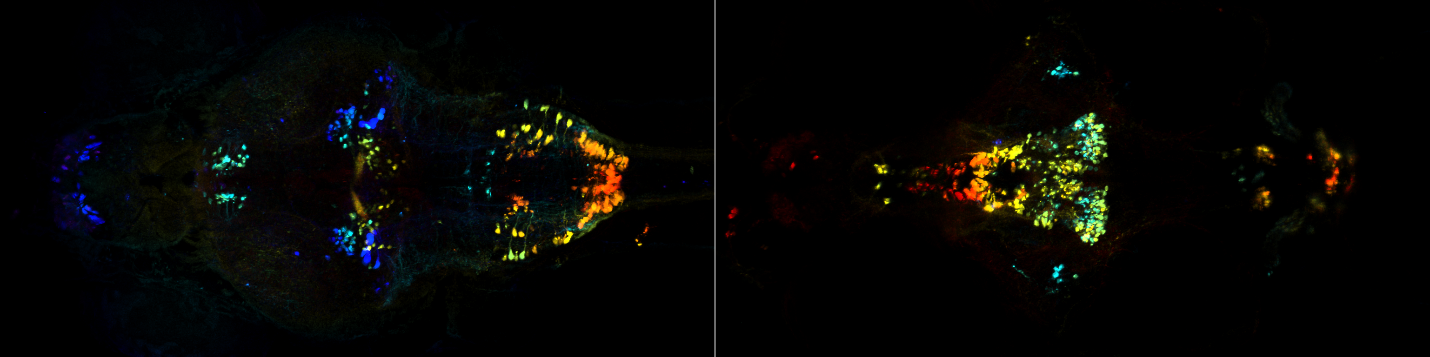

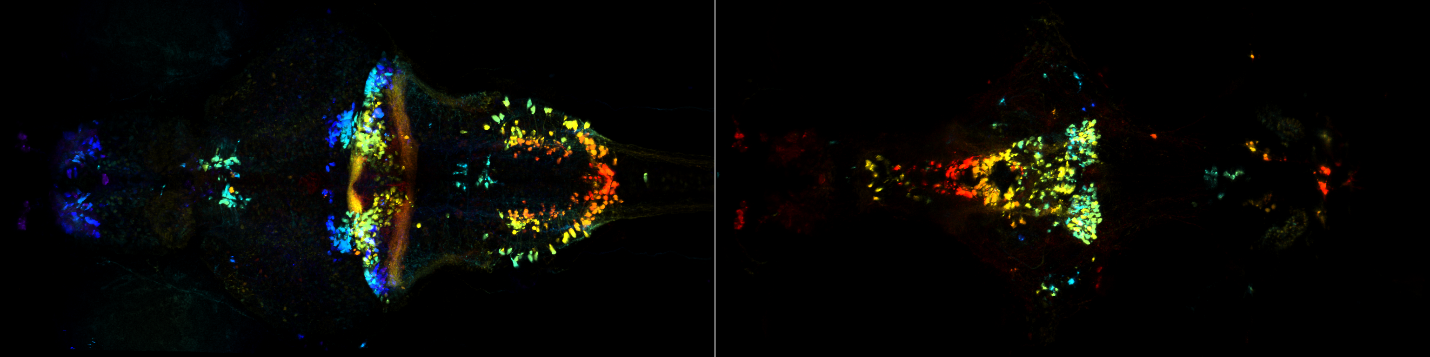

**Supplementary Fig. 2 | TH1 distribution in control.** Control Th1-Gal4 expression (depth-color-coded) in dorsal (left) and ventral (right) brain of 7-8 dpf larval zebrafish treated with 0.1% DMSO. Maximum projections were obtained across 200 µm for each half following registration using ANTs. Field-of-view: 0.5 mm (vertical) x 1.0 mm (horizontal).

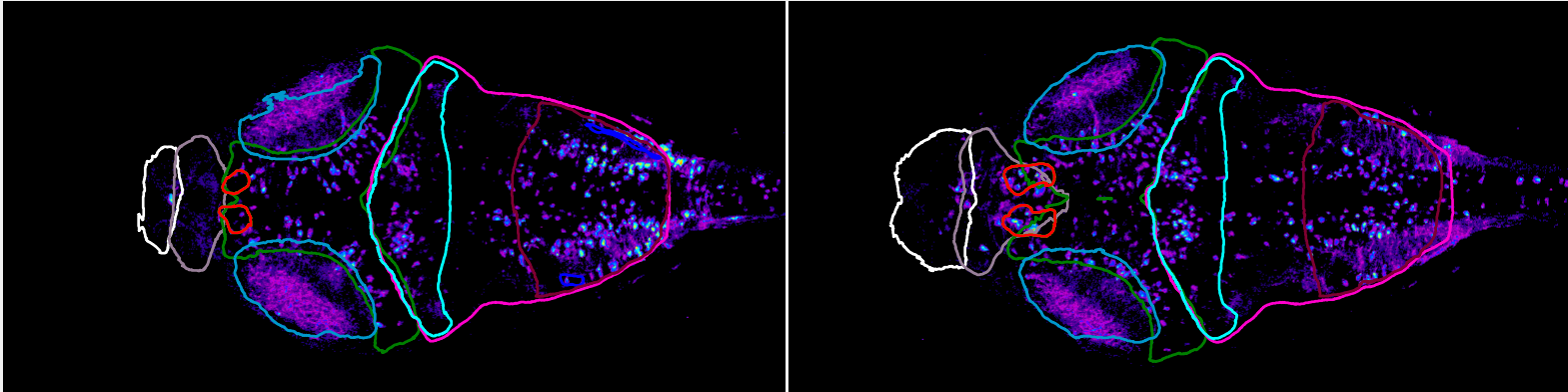

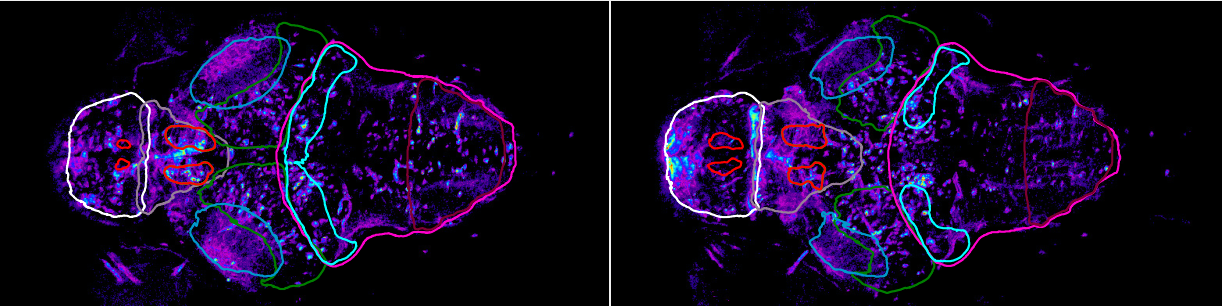

**Supplementary Fig. 3 | Mesencephalic TH1 and TH2 expression.** Th1- and Th2- expressing neurons can be seen inside #106 Mesencephalon Tectum Stratum Periventriculare (green) for four different 20-µm averaged Z-projections. Other anatomical masks shown for reference are #275 Telencephalon (white), #1 Diencephalon (grey), #107 Mesencephalon Tectum Neuropil (indigo), #114 Rhombencephalon (magenta), #131 Rho Cerebellum (cyan), and #225 Rhombomere 7 (cherry). Multiple regions delineated in red are dopaminergic clusters according to Z-brain atlas.

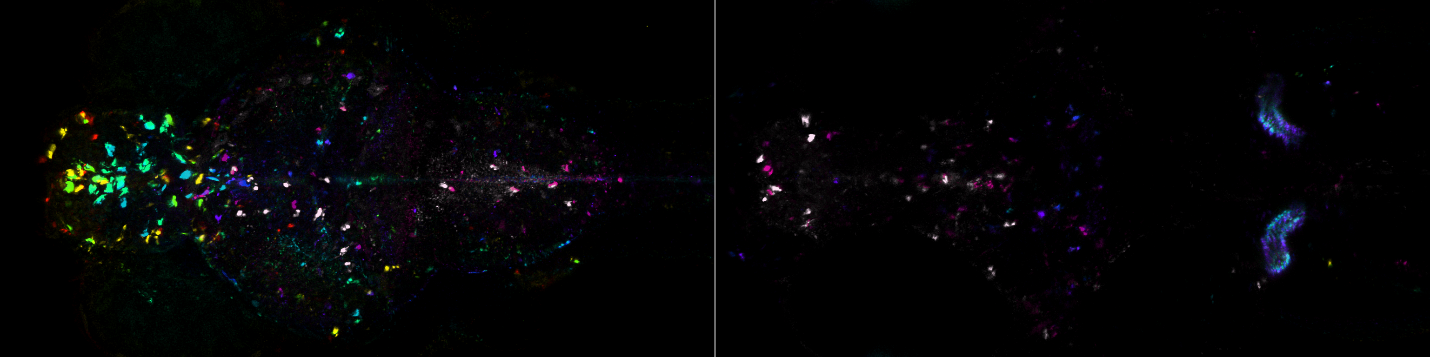

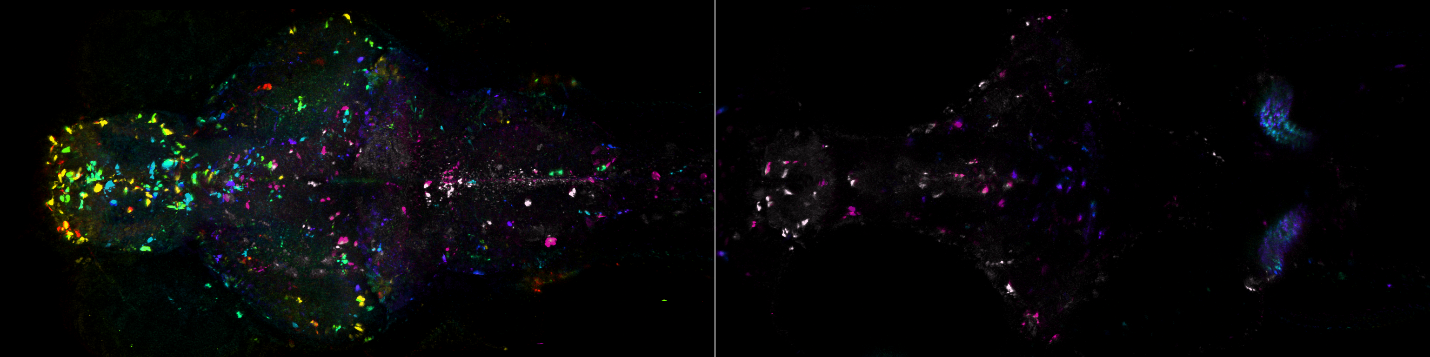

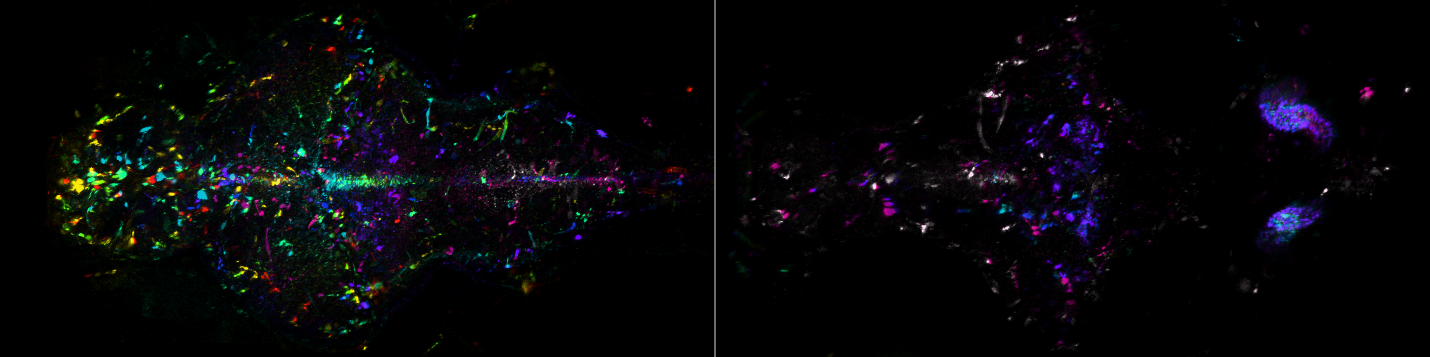

**Supplementary Fig. 4 | TH1 and TH2 distribution after chemogenetic ablation.** Depth-color-coded maximum projections of UAS-NTR-mCherry expression driven by*Th1-Gal4* and *Th2-Gal4* in dorsal (left) and ventral (right) brain of three 7-8 dpf larval zebrafish treated with 5 mM metronidazole (MTZ) in 0.1% DMSO for ~40 hours followed by 24-48 hours in system water. Maximum projections were obtained across 200 µm for each half following registration using ANTs. Field-of-view: 0.5 mm (vertical) x 1.0 mm (horizontal).

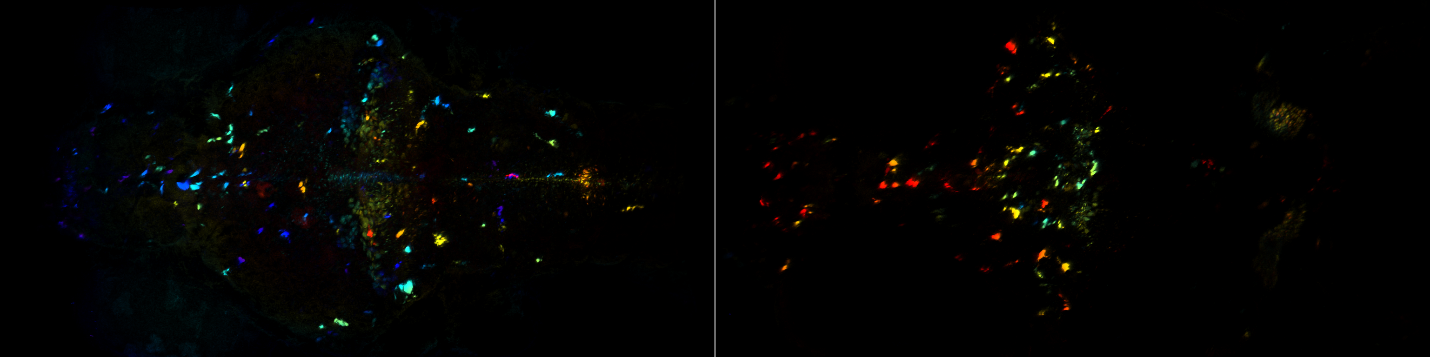

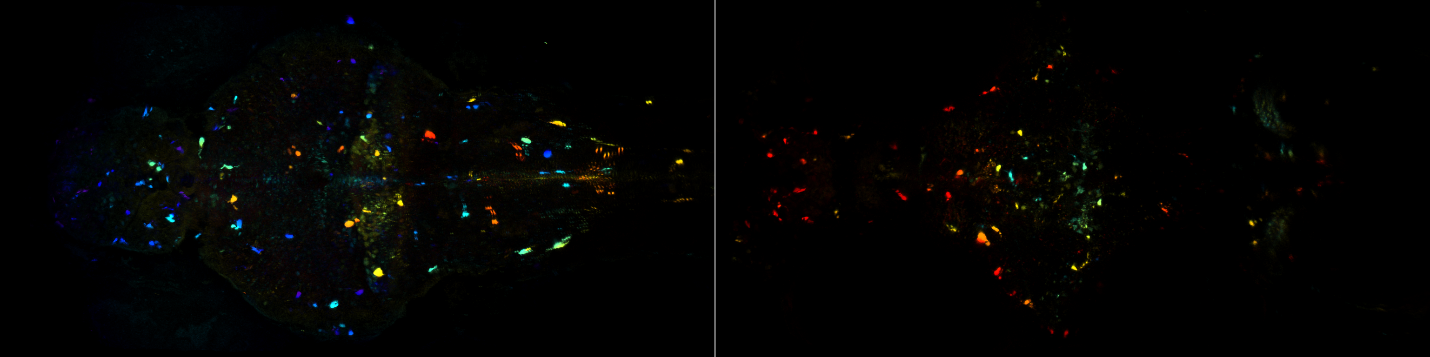

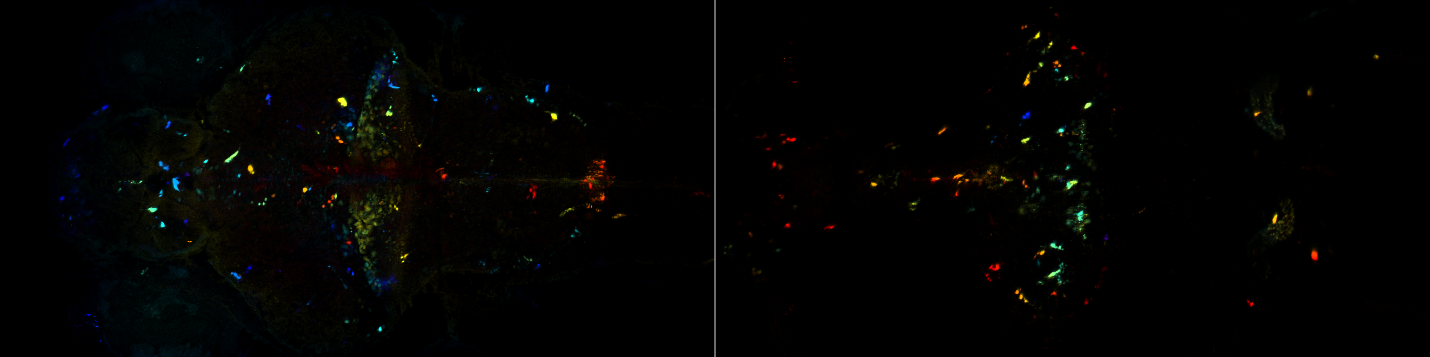

**Supplementary Fig. 5 | TH1 distribution following chemogenetic ablation.** Ablated Th1-Gal4 expression (depth-color-coded) in dorsal (left) and ventral (right) brain of 7-8 dpf larval zebrafish treated with 5 mM metronidazole (MTZ) in 0.1% DMSO. Maximum projections were obtained across 200 µm for each half following registration using ANTs. Field-of-view: 0.5 mm (vertical) x 1.0 mm (horizontal).

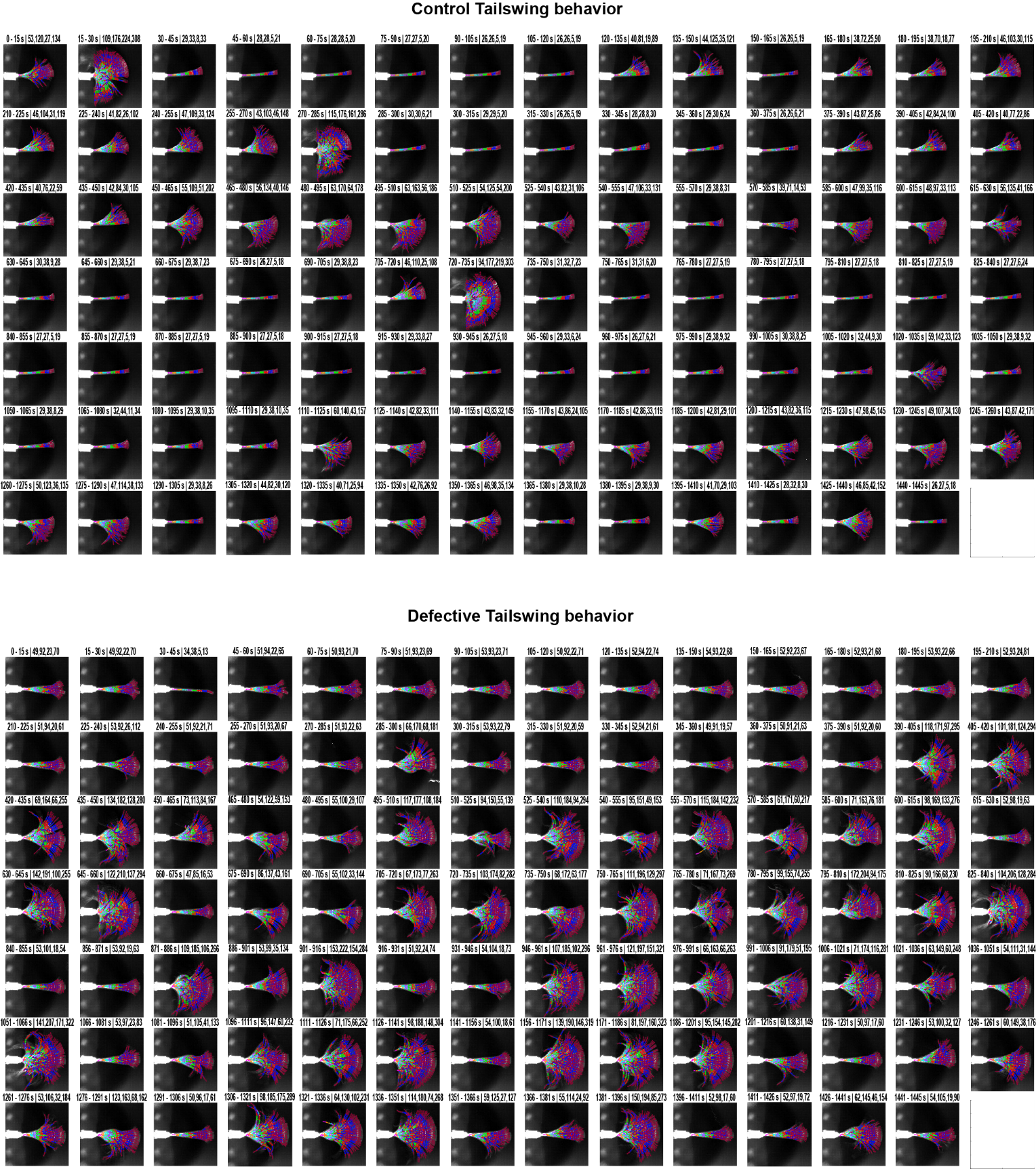

**Supplementary Fig. 6 | Automated tail tracking visualization using 15-second maximum projections.** Maximum projections of 15-second recordings overlaid with eight automatically detected tail segments (color-coded: magenta, orange, cyan, purple, green, red, blue, and cherry). Title parameters indicate: first two numbers - timestamps; third and fourth numbers - x-range (horizontal) for left and right tail halves; fifth and sixth numbers - y-range (vertical) for left and right tail halves, respectively, where higher right-half y-range values indicate greater tail activity. Tracking accuracy was validated by the close alignment of colored segments with the underlying tail image. Sub-optimal tracking can be improved by parameter adjustment.
